# Adaptive genomic compartments shaped by giant mobile elements underpin the ancient emergence of fungal pathogenicity

**DOI:** 10.64898/2026.05.11.723792

**Authors:** Andrea Menicucci, Chiara Fiorenzani, Firas Hatoum, Salvatore Iacono, Daniele Da Lio, Sioly Becerra, Gaétan Le Floch, Giovanni Vannacci, Aaron A. Vogan, Megan McDonald, Michael Thon, Serenella Sukno, Sabrina Sarrocco, Pedro Talhinhas, Riccardo Baroncelli

## Abstract

The emergence of new fungal pathogens often depends on the acquisition of complex adaptive traits, yet the mechanisms by which such traits arise remain poorly understood. Here we show that a biosynthetic gene cluster required for pathogenicity in the lupin pathogenic fungus *Colletotrichum lupini* was acquired within a genomic region derived from a giant *Starship* transposable element. Comparative and population genomic analyses reveal that the *C. lupini* genome contains multiple regions derived from ancestrally active *Starship* elements, enriched in lineage-specific genes and strongly induced during plant infection. One such region harbours a hybrid polyketide synthase-nonribosomal peptide synthetase (PKS-NRPS) gene cluster that is conserved in pathogenic isolates but absent from closely related non-pathogenic species and from a non-pathogenic strain. Phylogenetic analyses of the PKS-NRPS backbone gene reveal incongruence with species relationships and a distribution across deeply divergent fungal lineages, consistent with horizontal acquisition. Disruption of the PKS-NRPS backbone gene abolishes pathogenicity, demonstrating that this cluster is required for host infection. Phylogenomic analyses further indicate that lupin pathogenicity emerged once within the *C. lupini* lineage prior to its diversification. Together, these findings identify a *Starship*-associated virulence determinant and support a model in which giant cargo-mobilizing mobile elements generate genomic novelty by facilitating the acquisition, assembly and integration of adaptive traits during the emergence of fungal pathogenicity.

## INTRODUCTION

Fungal plant pathogens are a major constraint on global crop production, causing substantial losses in yield and quality and posing an increasing threat to food security^1–4^. The emergence of new diseases is often driven by rapid evolutionary transitions, including host shifts and the acquisition of novel virulence traits, which enable pathogens to colonize previously inaccessible hosts^5–9^. Understanding the genetic mechanisms that underlie these transitions is therefore critical for predicting pathogen emergence and developing sustainable control strategies.

Host specialization and host shifts are frequently mediated by genes involved in host recognition, immune suppression and nutrient acquisition, including secreted effectors, detoxification enzymes, carbohydrate-active enzymes and secondary metabolite biosynthetic clusters^10–12^. Increasing evidence indicates that genome plasticity plays a central role in generating such adaptive innovations. Horizontal gene transfer, accessory chromosomes and transposable elements can introduce novel genetic material into pathogen genomes, accelerating adaptation to new hosts and environments^10,13–21^. However, how complex, multi-gene traits are assembled, integrated and retained within fungal genomes remains poorly understood.

Recently identified giant cargo-mobilizing mobile elements have added a new dimension to this problem. Among them, *Starship* elements represent a class of exceptionally large transposable elements encoding *Captain* tyrosine recombinase (DUF3435 domain) proteins capable of mobilizing extensive genomic regions containing functional genes^22,23^. These elements are widespread across fungal genomes and have been proposed to mediate horizontal transfer of adaptive traits at large scales^24^. Although *Starships* capacity for genome-scale mobility suggests a potentially important role in the emergence of complex traits, their long-term evolutionary impact remains unclear, particularly in cases where mobility has been lost and only remnant sequences persist.

The genus *Colletotrichum* comprises numerous plant-associated fungi responsible for economically important diseases across a wide range of crops^25–29^. Within this genus, anthracnose of lupins (*Lupinus* spp.) is exclusively caused by a single species, *Colletotrichum lupini*^30,31^. This striking host specificity suggests that the ability to infect lupin arose through a discrete evolutionary transition. However, the timing, origin and genetic basis of this adaptation remain unknown.

Here, we investigate the emergence of lupin pathogenicity in *C. lupini* using an integrative framework combining comparative genomics, population genomics, transcriptomics and functional genetics. We identify multiple genomic regions derived from giant cargo-mobilizing mobile elements that define dynamic compartments in the genome and carry genes that are strongly induced during plant infection. Although these elements lack conserved catalytic features required for mobility, indicating that they are degenerated remnants of ancestrally active elements, they retain functional relevance. Within one such region, we identify a single horizontally transferred hybrid polyketide synthase-nonribosomal peptide synthetase (PKS-NRPS) biosynthetic gene cluster that is conserved among pathogenic isolates and required for host infection. Together, our results support a model in which giant cargo-mobilizing mobile elements facilitated the assembly and integration of a key adaptive trait during the emergence of pathogenicity, and whose degenerated remnants now persist as functionally important genomic compartments.

## RESULTS

### Lupin pathogenicity originated once within the *Colletotrichum lupini* lineage

To investigate the evolutionary origin of lupin pathogenicity, we combined phylogenomic reconstruction with experimental infection assays across representative *Colletotrichum* species and *C. lupini* isolates (Fig. 1A & B; Supplementary file 1). Pathogenicity assays on *Lupinus albus* revealed a clear dichotomy between *C. lupini* and closely related species: all tested *C. lupini* isolates induced characteristic anthracnose symptoms, including stem twisting, necrotic lesions and plant collapse, whereas non-*C. lupini* species failed to cause disease under identical conditions (Fig. 1B). This pattern was consistent across seedling and detached leaf assays, indicating that the ability to infect lupin is largely restricted to *C. lupini*. Notably, pathogenicity did not correlate with host of isolation, as the *C. lupini* strain CBS129944, originally isolated from *Cinnamomum burmanni*, was fully pathogenic on lupin, whereas the distantly related species *C. tofieldiae*, isolated from *Lupinus polyphyllus*, did not induce symptoms. Within *C. lupini*, all isolates were pathogenic except strain JA20 (lineage 3), indicating that pathogenicity is largely conserved at the species level but may vary among lineages.

**Fig. 1.**
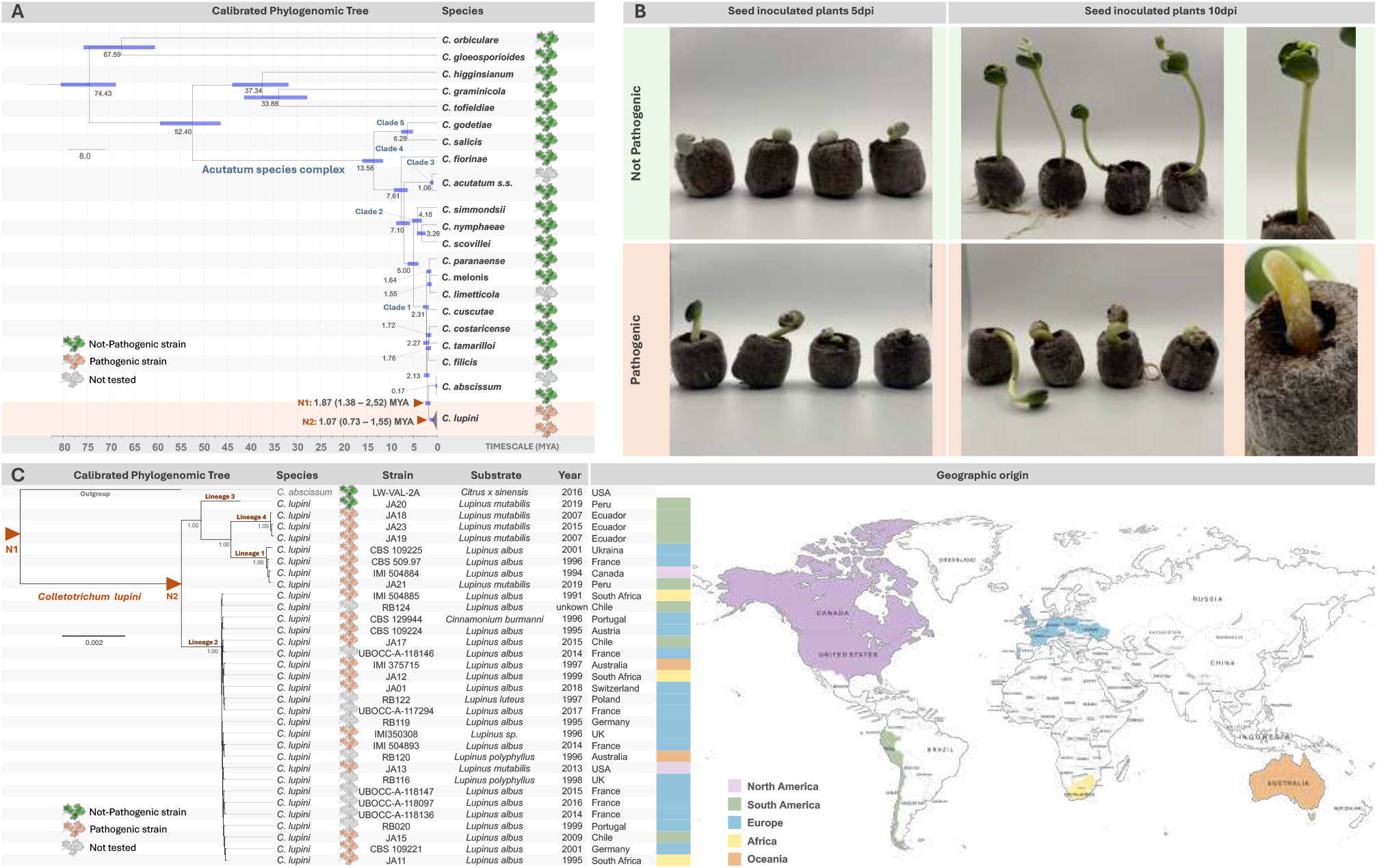
Evolutionary origin of lupin pathogenicity and phylogenomic diversity of *Colletotrichum lupini*. **A**, Time-calibrated phylogeny of representative *Colletotrichum* species inferred from genome-wide single-copy orthologs, with *Verticillium dahliae* used as outgroup. Branch lengths indicate divergence times estimated using the RelTime method and are scaled in million years ago (MYA). Blue bars at nodes represent 95% confidence intervals for divergence-time estimates. Major clades within the Acutatum species complex are indicated, and the position of *C. lupini* is highlighted. Symbols adjacent to species names denote pathogenic, non-pathogenic or untested strains in lupin pathogenicity assays. Estimated divergence times for nodes N1 and N2 are indicated. **B**, Pathogenicity assays on *Lupinus albus* seedlings inoculated with representative *Colletotrichum* strains. Images show seedlings at 5 and 10 days post inoculation (dpi). Non-pathogenic strains did not induce disease symptoms, whereas pathogenic *C. lupini* strains caused characteristic anthracnose symptoms, including seedling collapse, hypocotyl twisting and tissue necrosis. Representative close-up images of symptomatic seedlings are shown on the right. **C**, Phylogenomic relationships among *C. lupini* isolates inferred from genome-wide single-copy orthologs. Isolates are annotated by lineage, host or substrate of origin, year of isolation and geographic origin. Four major *C. lupini* lineages are identified, with *C. abscissum* included as outgroup. Symbols indicate pathogenic, non-pathogenic or untested strains. Coloured bars denote geographic origin, and the accompanying world map summarizes the global distribution of sampled isolates across Europe, North America, South America, Africa and Oceania.

To place these observations in an evolutionary context, we reconstructed a time-calibrated tree using a dataset derived from single-copy orthologues across the full proteomes (Fig. 1A). All pathogenic isolates clustered within a single monophyletic lineage corresponding to *C. lupini*, whereas non-pathogenic taxa were distributed across the broader *Colletotrichum* phylogeny. This topology is consistent with a scenario in which the ability to infect lupin originated once in the common ancestor of *C. lupini*, rather than arising through multiple independent events. Divergence time estimates indicate that *C. lupini* split from its closest relative, *C. abscissum*, approximately 1.87 million years ago (95% HPD: 1.38-2.52 MYA), with subsequent diversification beginning around 1.07 MYA (95% HPD: 0.73-1.55 MYA), predating modern agriculture and suggesting that the emergence of pathogenicity occurred in a natural ecological context.

Phylogenomic analysis further revealed a strongly structured population subdivided into four distinct lineages with clear geographic associations (Fig. 1C). The highest genetic diversity was observed in South America, particularly in the Andean region, where multiple lineages coexist. In contrast, a single lineage (lineage II) has undergone recent global expansion and underlies the contemporary lupin anthracnose pandemic. The seed-borne nature of *C. lupini* likely facilitated long-distance dispersal via international trade. Together, these results indicate that lupin pathogenicity is a species-associated trait that likely emerged prior to global dissemination and has been broadly maintained across lineages.

### Lineage-specific genome remodeling concentrates gene gain, loss and fragmentation in dynamic regions of *Colletotrichum lupini*

To investigate the genomic basis of host specialization, we compared high-quality genomes of *C. lupini* with those of closely related species (Fig. 2A). This analysis revealed extensive lineage-specific remodeling of gene content, including both gene loss and gene gain events concentrated in discrete genomic regions.

**Fig. 2.**
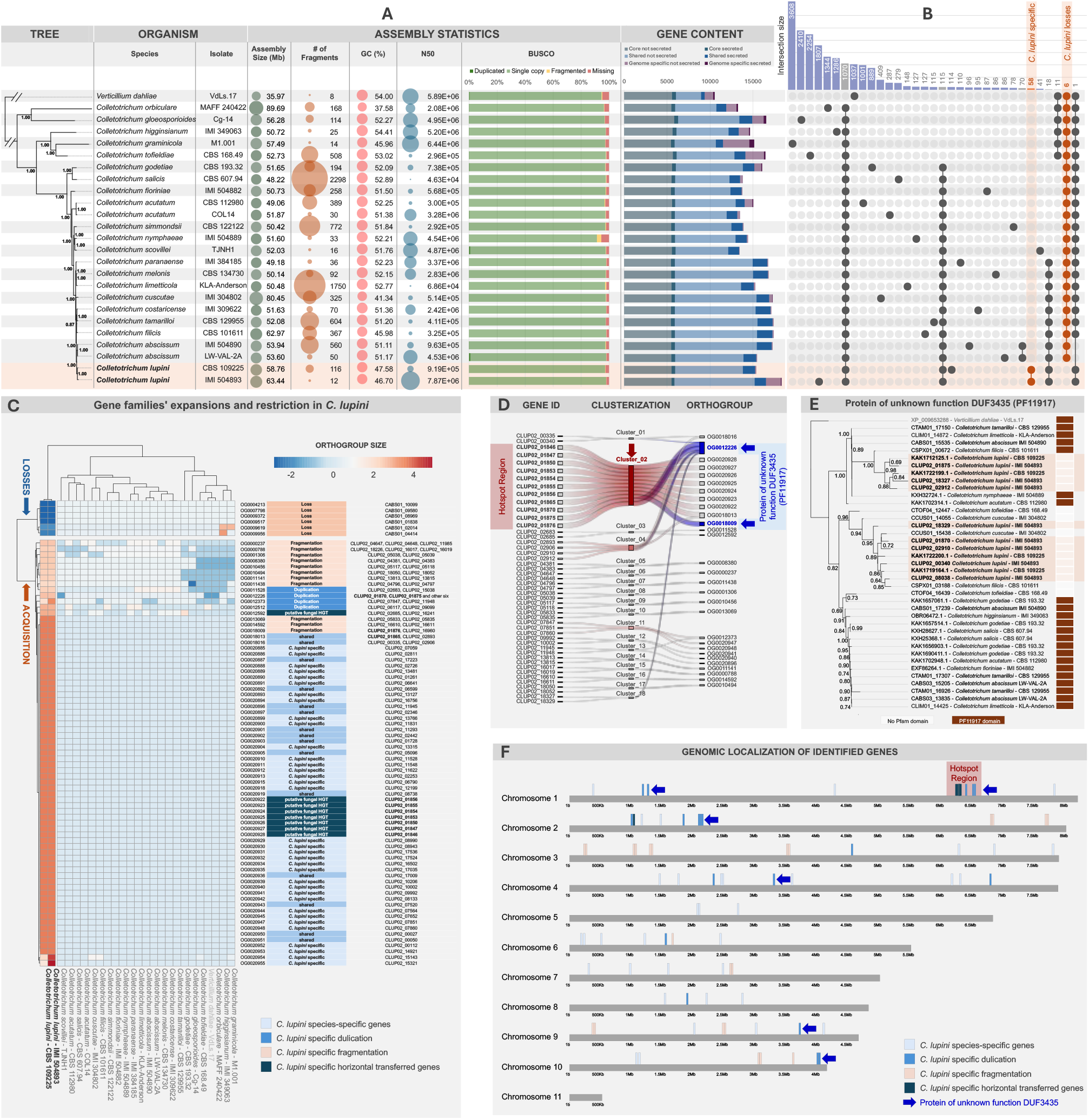
Lineage-specific genome remodelling and dynamic genomic regions in *C. lupini*. **A**, Phylogenomic relationships and genome features of the analysed *Colletotrichum* species and the outgroup *Verticillium dahliae*. Bubble plots show genome size, GC content, assembly contiguity (N50) and predicted gene content. Gene categories are partitioned into core genes shared by all analysed species, shared genes present in at least two but not all species, and species-specific genes; secreted and non-secreted subsets are indicated by colour. Bubble size is proportional to gene number. **B**, Orthogroup presence-absence patterns across species highlighting lineage-specific gene gain and loss in *C. lupini*. Vertical bars indicate orthogroup intersection size, whereas connected dots indicate species combinations sharing each orthogroup. Orange labels denote orthogroups associated with *C. lupini*-specific acquisitions or losses. **C**, Gene-family expansions and contractions in *C. lupini*. Heatmaps show relative enrichment or depletion of orthogroup copy number compared with related species, with blue and red colours indicating contractions and expansions, respectively. Orthogroups associated with species-specific genes, duplications, fragmentations and putative horizontal gene transfer events are highlighted. **D**, Clustering of lineage-specific, duplicated, fragmented and putatively horizontally acquired genes identifies discrete dynamic genomic regions (“hotspots”). Coloured links connect genes belonging to the same orthogroup, whereas labelled clusters denote hotspot regions enriched in these features. **E**, Phylogenetic relationships of DUF3435-domain proteins, homologous to tyrosine site-specific recombinases, identified across multiple loci and *Colletotrichum* genomes. Branch labels correspond to protein identifiers and genome origins. Brown boxes indicate the conserved PF11917-related DUF3435 domain, whereas white boxes indicate the absence of a detectable domain. Numbers at nodes indicate branch support values. **F**, Genome-wide distribution of species-specific genes, duplicated genes, fragmented genes, putatively horizontally acquired genes and DUF3435-containing genes across the 11 chromosomes of *C. lupini*. Highlighted regions indicate hotspots enriched in dynamic genomic features.

We identified six orthogroups conserved across related species but entirely absent in *C. lupini*, including genes encoding a predicted transcription factor and a RING-type E3 ubiquitin ligase, suggesting loss of regulatory functions (Fig. 2B; Supplementary file 2 and 3). Conversely, 58 orthogroups were initially classified as specific to *C. lupini*, with a subset showing expansion relative to close relatives (Fig. 2C). However, expanded taxon sampling indicated that some of these genes are present in other *Colletotrichum* species, highlighting the sensitivity of lineage-specific inferences to taxon sampling.

Strikingly, most expanded orthogroups were not associated with canonical gene duplication but instead with insertion of highly similar 5-6 kb transposable elements, resulting in gene fragmentation (Fig. 2C; Supplementary file 4). These fragments show high sequence conservation across loci and are frequently associated with disrupted gene structures lacking detectable expression, indicating that insertion-mediated inactivation contributes substantially to genome remodeling in *C. lupini*. Only a minority of expansions represent whole gene duplications, including genes encoding Peptidase M49 family proteins, some of which display phylogenetic relationships inconsistent with the species tree and may reflect more complex evolutionary histories.

Genes gained, expanded or fragmented in *C. lupini* are not randomly distributed but instead cluster in discrete genomic regions. Notably, one hotspot region encompassing several genes with locus tags beginning with CLUP02_018 was enriched in genes belonging to multiple orthogroups, including several lineage-restricted or phylogenetically atypical genes (Fig. 2D & F). Phylogenetic analyses indicate that some of these genes share related evolutionary patterns, consistent with their accumulation within linked genomic regions followed by local rearrangement and fragmentation.

A recurrent feature of these regions is the presence of genes encoding DUF3435-domain proteins, homologous to tyrosine site-specific recombinases (Fig. 2E).

These genes are distributed across multiple loci and frequently co-occur with duplicated orthogroups, suggesting a shared evolutionary origin linked to mobile genetic elements and ancient duplication events in the ancestor of *C. lupini*. However, the absence of conserved catalytic residues in all identified proteins indicates loss of recombinase activity, consistent with functional degeneration. Together, these observations support a model in which these regions represent derelict *Starships*, namely remnants of ancestrally active, giant cargo-mobilizing mobile elements that have undergone inactivation, loss of mobility, and progressive genomic decay.

These results indicate that genome evolution in *C. lupini* is shaped by the accumulation and progressive degeneration of *Starships*, generating discrete genomic compartments enriched in gene gain, loss and insertion-mediated fragmentation. Although several genes within these regions phylogenetic relationships that deviate from the expected species lineage, evidence for horizontal acquisition is most clearly supported for the biosynthetic gene cluster described below.

### Derelict *Starships* define dynamic genomic compartments enriched in infection-induced genes

Comparative genomic analyses identified multiple highly dynamic regions in the *C. lupini* genome that are enriched in recombinase-related genes. Macrosynteny comparisons with closely related species showed that overall chromosome organization is largely conserved, but several discrete regions are present in *C. lupini* and absent from its relatives, indicating localized structural variation rather than pervasive chromosomal reshuffling (Fig. 3A). Microsynteny analyses further showed that these regions occur as large insertions embedded within otherwise collinear chromosomes.

**Fig. 3.**
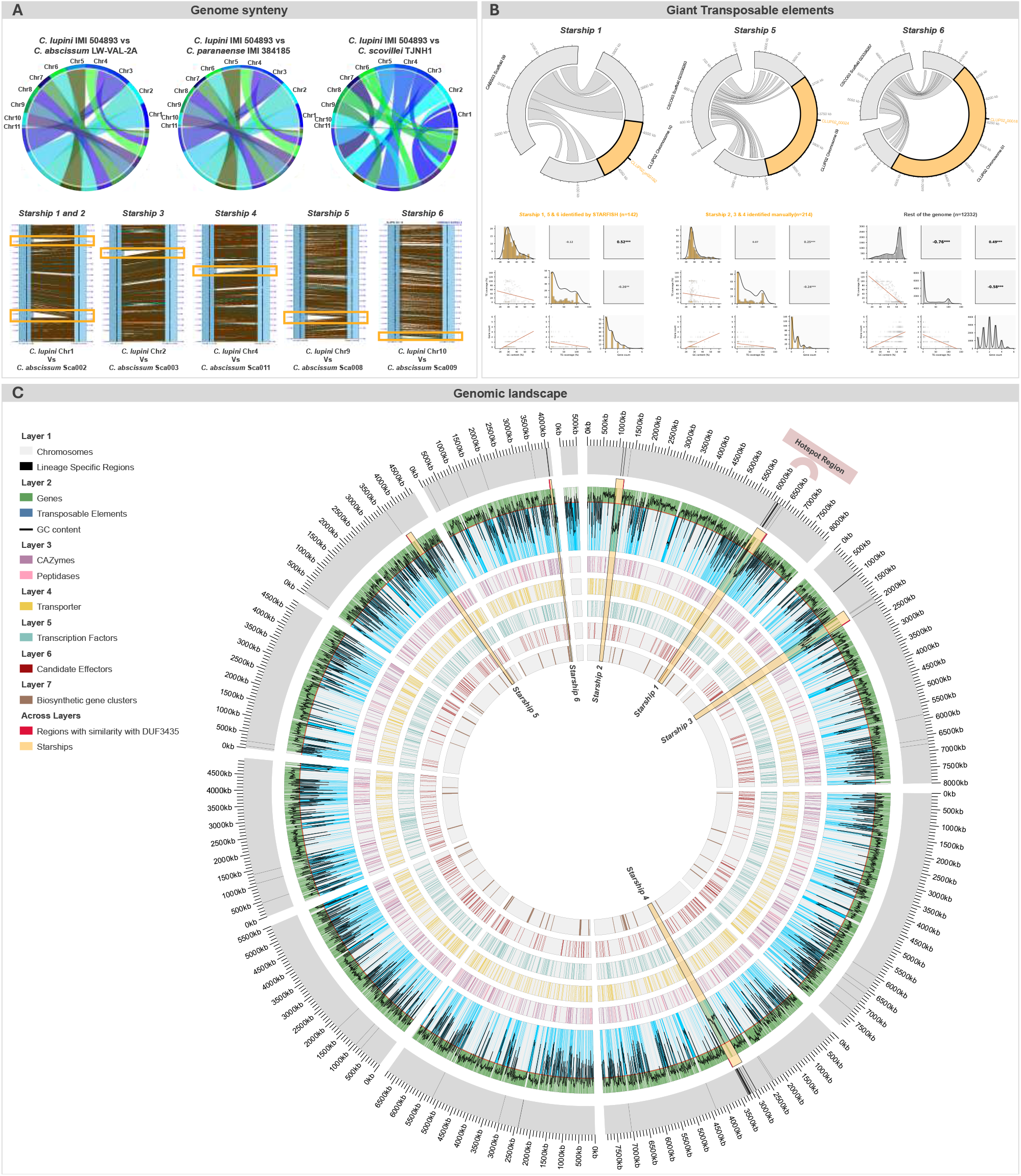
Derelict *Starships* define dynamic genomic compartments in *C. lupini*. **A**, Macro- and microsynteny comparisons between *C. lupini* and closely related species. Top, whole-chromosome synteny relationships between *C. lupini* IMI 504893 and the related species *C. abscissum* LW-VAL-2A, *C. paranaense* IMI 384185 and *C. scovillei* TJNH1. Ribbons connect syntenic genomic regions across chromosomes. Bottom, detailed microsynteny analyses of representative insertion loci corresponding to *Starships* 1-6, highlighting lineage-specific insertions in *C. lupini* relative to homologous regions in related species. Orange boxes indicate insertion boundaries. **B**, Identification and structural characterization of giant mobile elements derived regions (derelict *Starships*). Top, detection of insertion sites through comparison with corresponding empty loci in related species. Bottom, structural comparisons and feature distributions for intact and degenerated *Starship* regions identified using automated (STARFISH) and manual approaches. Density plots compare genomic features associated with *Starships* 1, 5 and 6 identified by STARFISH, *Starships* 2, 3 and 4 identified manually, and the remainder of the genome. **C**, Genome-wide distribution of derelict *Starships* and associated genomic features. Circos plot showing the 11 chromosomes of *C. lupini* and lineage-specific regions together with tracks representing gene density, transposable element density and GC content. Additional layers show the distribution of carbohydrate-active enzymes (CAZymes), peptidases, transporters, transcription factors, candidate effectors and biosynthetic gene clusters. Inner tracks indicate regions with similarity to DUF3435-containing recombinases and the localization of *Starship* regions. Highlighted sectors denote genomic hotspots enriched in dynamic genomic features.

These dynamic regions overlap with loci identified in the comparative genomic analysis, including the region previously designated as a hotspot. More broadly, these regions consistently contain genes encoding DUF3435-domain proteins. The recurrent association between structurally distinct regions, phylogenetically atypical genes and recombinase-related domains is consistent with their origin from giant cargo-mobilizing mobile elements. To systematically characterize these regions, we combined automated detection using Starfish with manual curation of additional loci sharing similar structural and compositional features (Fig. 3B). Together, these analyses identified six regions distributed across multiple chromosomes. Although some retain partial structural hallmarks consistent with previously described cargo-carrying elements, all lack key features required for mobility, including intact recombinase catalytic domains.

These observations suggest that the identified regions correspond to derelict *Starships*, representing degenerated remnants of ancestrally active mobile elements that have likely lost their capacity for mobilization.

Genome-wide mapping showed that these regions define a distinct genomic compartment enriched in lineage-specific sequences, transposable elements and genes linked to host interaction, including biosynthetic gene clusters (Fig. 3C). Transcriptomic analyses further revealed that genes within these regions are preferentially induced during plant infection compared with *in vitro* conditions (Fig. 4; Supplementary file 5), with several among the most strongly upregulated genes *in planta*.

**Fig. 4.**
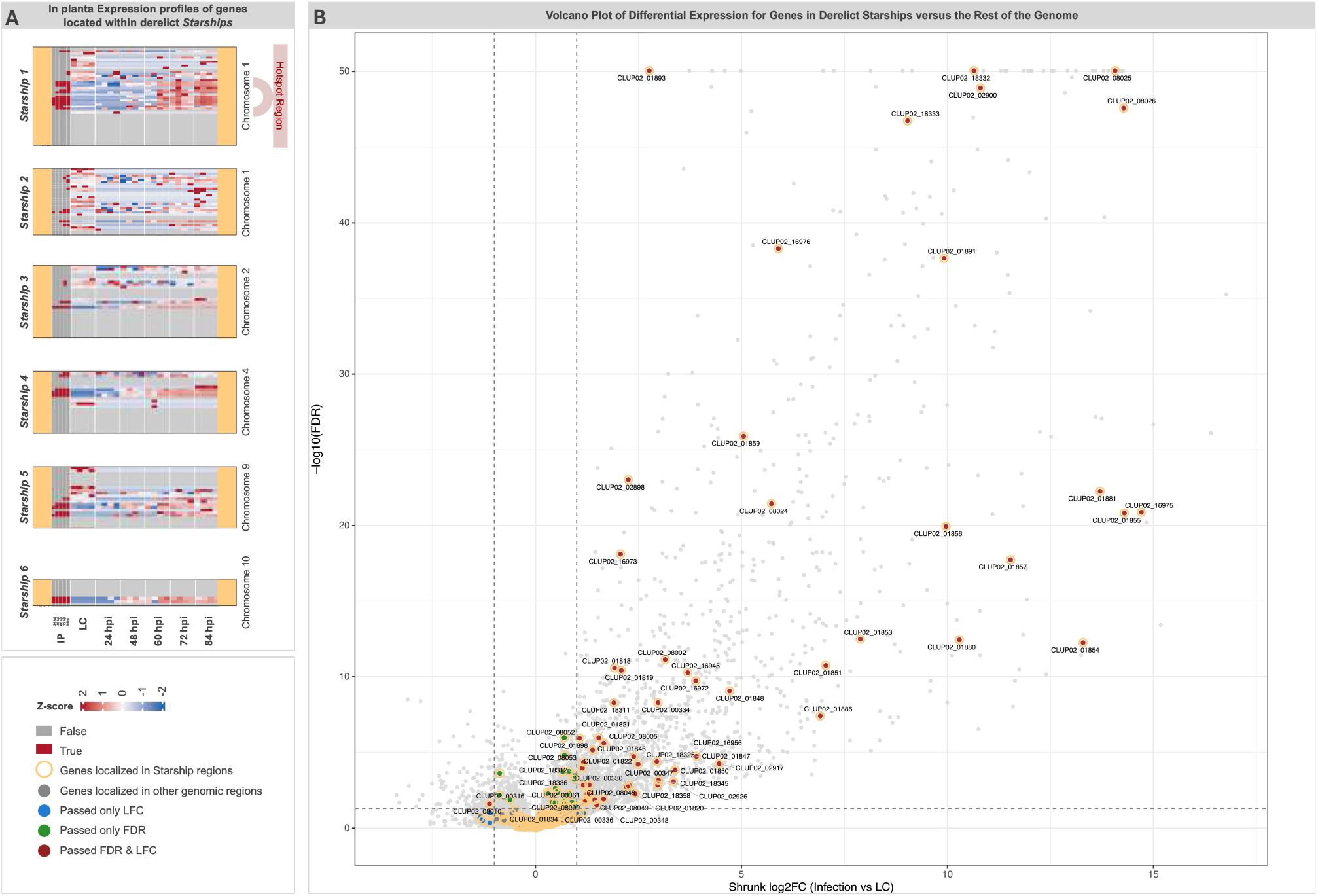
Derelict *Starships* are characterized by highly infection-induced genes during plant infection. **A**, Expression profiles of genes located within derelict *Starship* regions across *in vitro* and *in planta* conditions. Heatmaps show row z-score-normalized gene expression values for genes located within Starships 1-6 across liquid culture (LC) and infected plant tissue sampled at 24, 48, 60, 72 and 84 hours post inoculation (hpi). Columns represent individual biological replicates for each condition, whereas rows represent genes ordered according to their genomic position within each *Starship* region. Red and blue colours indicate relative upregulation and downregulation, respectively. Hotspot-associated region corresponding to *Starship* 1 is indicated. **B**, Differential expression analysis of genes located within derelict *Starships* compared with genes located in the remainder of the genome during infection. Volcano plot showing log2 fold change in expression between in planta and liquid culture conditions versus −log10 adjusted *P* value (FDR). Genes located within derelict *Starships* are highlighted and labelled, whereas genes located elsewhere in the genome are shown in grey. Vertical and horizontal dashed lines indicate fold-change and significance thresholds, respectively. Points are coloured according to significance criteria, including genes passing fold-change threshold only, FDR threshold only, or both thresholds simultaneously.

Together, these results indicate that derelict *Starships* represent structurally and transcriptionally distinct genomic compartments that are associated with infection-related gene expression. While several genes within these regions show atypical evolutionary patterns, robust evidence for horizontal acquisition is restricted to some genes within the biosynthetic gene cluster described below.

### A derelict *Starship*-associated biosynthetic gene cluster is conserved in pathogenic isolates and induced during infection

Within the derelict *Starships* identified in *C. lupini*, one locus corresponding to the hotspot region on chromosome 1 (*Starship 1*) showed multiple features consistent with a role in pathogenicity. This region is present in *C. lupini* but absent from the closely related species *C. abscissum*, indicating lineage-specific acquisition (Fig. 5A). We therefore focused on *Starship 1* to investigate whether these derelict *Starships* encode candidate pathogenicity determinants.

**Fig. 5.**
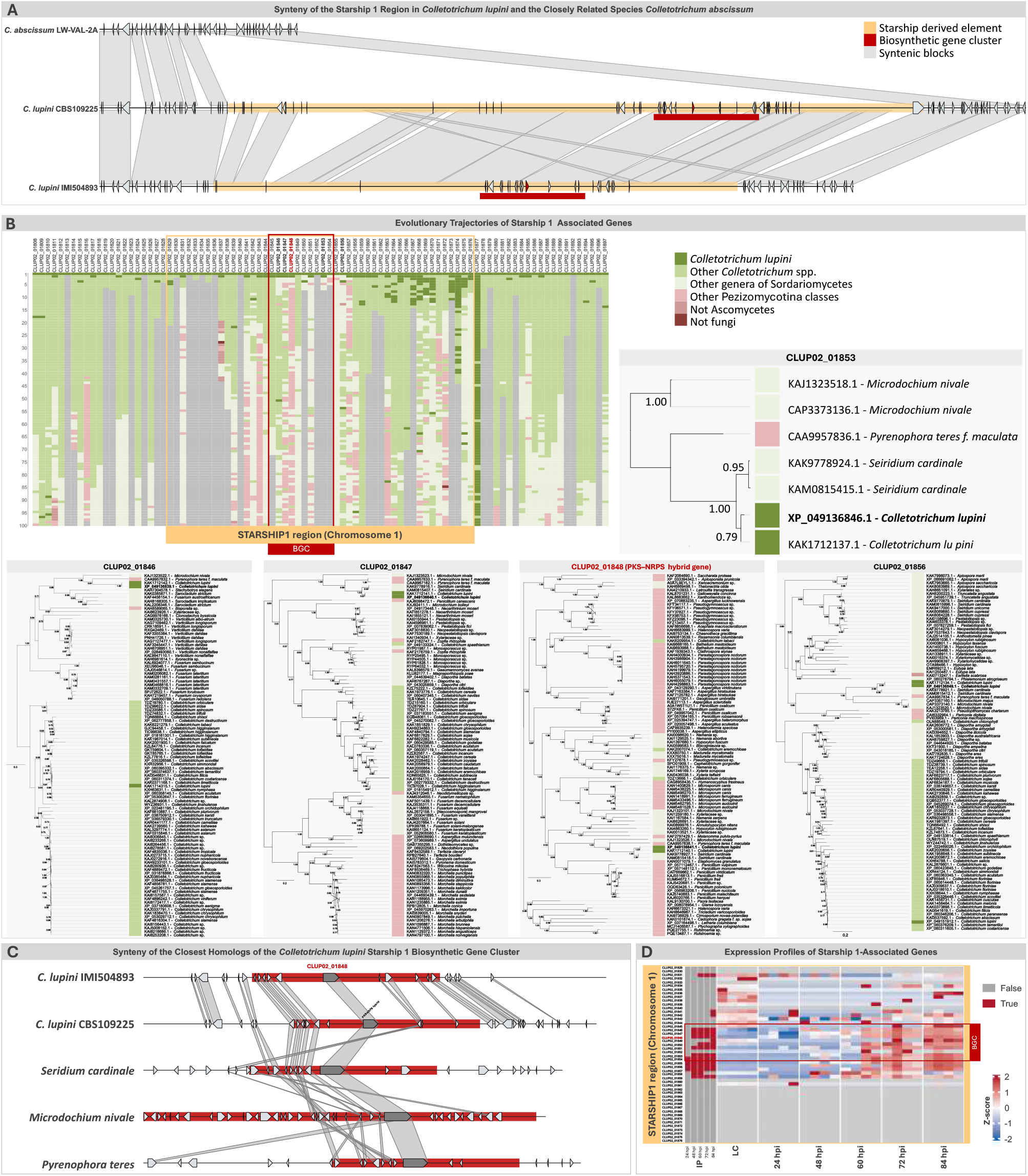
A derelict Starship harbours a horizontally acquired PKS-NRPS biosynthetic gene cluster linked to pathogenicity. **A**, Presence-absence comparison of derelict *Starship* 1 region between *C. lupini* and the closely related species *C. abscissum*. Synteny analyses reveal a lineage-specific insertion in *C. lupini* containing a biosynthetic gene cluster (BGC) absent from the homologous region in *C. abscissum*. Grey ribbons indicate syntenic blocks between genomes, whereas orange regions denote derelict *Starship* 1 and red regions indicate the BGC. **B**, Evolutionary trajectories of genes associated with the derelict *Starship* 1 region. Top, taxonomic distribution of the top 100 most similar proteins identified for genes located across the *Starship 1* region, revealing heterogeneous phylogenetic affinities and a mosaic evolutionary origin. Colours indicate homologues from *C. lupini*, other *Colletotrichum* species, other Sordariomycetes, other Pezizomycotina classes, non-fungal taxa and non-ascomycete lineages. Bottom, phylogenetic analyses of representative genes from the hotspot region, including the hybrid polyketide synthase-nonribosomal peptide synthetase (PKS-NRPS) backbone gene CLUP02_01848 and neighbouring genes. *C. lupini* sequences do not cluster with homologues from closely related *Colletotrichum* species, but instead group with distantly related fungi from multiple fungal classes, supporting horizontal acquisition of the cluster or some of its components. Node labels indicate branch support values. **C**, Gene content and structural organization of the derelict *Starship* 1 hotspot region and comparison with the closest homologous regions identified in other fungi, including *Seiridium cardinale, Microdochium nivale* and *Pyrenophora teres*. Several genes show highest similarity to distantly related fungal taxa, consistent with a mosaic evolutionary origin of the region. Syntenic blocks connecting homologous genes are shown in grey. **D**, Expression profiles of genes located within the derelict *Starship* 1region during infection. Heatmaps show row z-score-normalized expression values across liquid culture (LC) and infected plant tissue sampled at 24, 48, 60, 72 and 84 hours post inoculation (hpi). Genes belonging to the PKS-NRPS biosynthetic gene cluster are strongly induced in planta compared with in vitro conditions. Columns represent individual biological replicates, and rows represent genes ordered according to genomic position within the region

The region spans approximately 400 kb and contains a mixture of genes, including lineage-restricted genes lacking detectable homologues outside the genus and genes with conserved domains showing similarity to distantly related fungi. Among these, a cluster of genes (CLUP02_01842-CLUP02_01855) forms a predicted hybrid polyketide synthase-nonribosomal peptide synthetase (PKS-NRPS) biosynthetic gene cluster (Fig. 5B). The cluster encodes a complete set of biosynthetic and accessory functions, including a PKS-NRPS backbone enzyme, cytochrome P450 monooxygenase, oxidoreductases and putative regulatory components.

Comparative analyses revealed that this cluster has a heterogeneous similarity profile (Fig. 5B). While some genes are most similar to homologues within *Colletotrichum*, others show highest similarity to genes from distantly related fungi, including *Seiridium cardinale, Microdochium nivale*, and *Pyrenophora teres* (Fig. 5B). Phylogenetic analysis of the PKS-NRPS backbone gene also showed incongruence with the species phylogeny, as *C. lupini* sequences clustered with distantly related fungi rather than with closely related *Colletotrichum* species (Fig. 5B). In addition, homologues of this cluster are distributed across diverse fungal lineages, suggesting a complex evolutionary history not explained by vertical inheritance alone. The BGC displays high synteny and sequence similarity across these distantly related fungi. Together with its absence from closely related species (Fig. 5C), this pattern is consistent with horizontal acquisition of the cluster or its components. The broad distribution of homologues across distant fungal taxa further suggests horizontal mobility of these genes, potentially involving multiple transfer events or ancient dissemination.

Consistent with a functional role during host interaction, transcriptomic analyses showed that genes within this region are strongly induced during plant infection. The PKS-NRPS cluster, in particular, is among the most highly expressed loci *in planta* across multiple infection time points, contrasting with low expression under in vitro conditions (Fig. 5D). Together, these features identify this derelict *Starship*-associated biosynthetic gene cluster as a strong candidate determinant of pathogenicity in *C. lupini*.

### The PKS-NRPS backbone gene is required for pathogenicity

To test whether the derelict *Starship*-associated biosynthetic gene cluster contributes to pathogenicity, we disrupted the PKS-NRPS backbone gene in the reference strain IMI504893 and assessed infection phenotypes (Fig. 6). Deletion of the PKS-NRPS gene (CLUP02_01848) resulted in complete loss of pathogenicity: while the wild-type strain produced typical anthracnose symptoms on lupin tissues, both independent deletion mutants failed to induce disease under identical conditions.

**Fig. 6.**
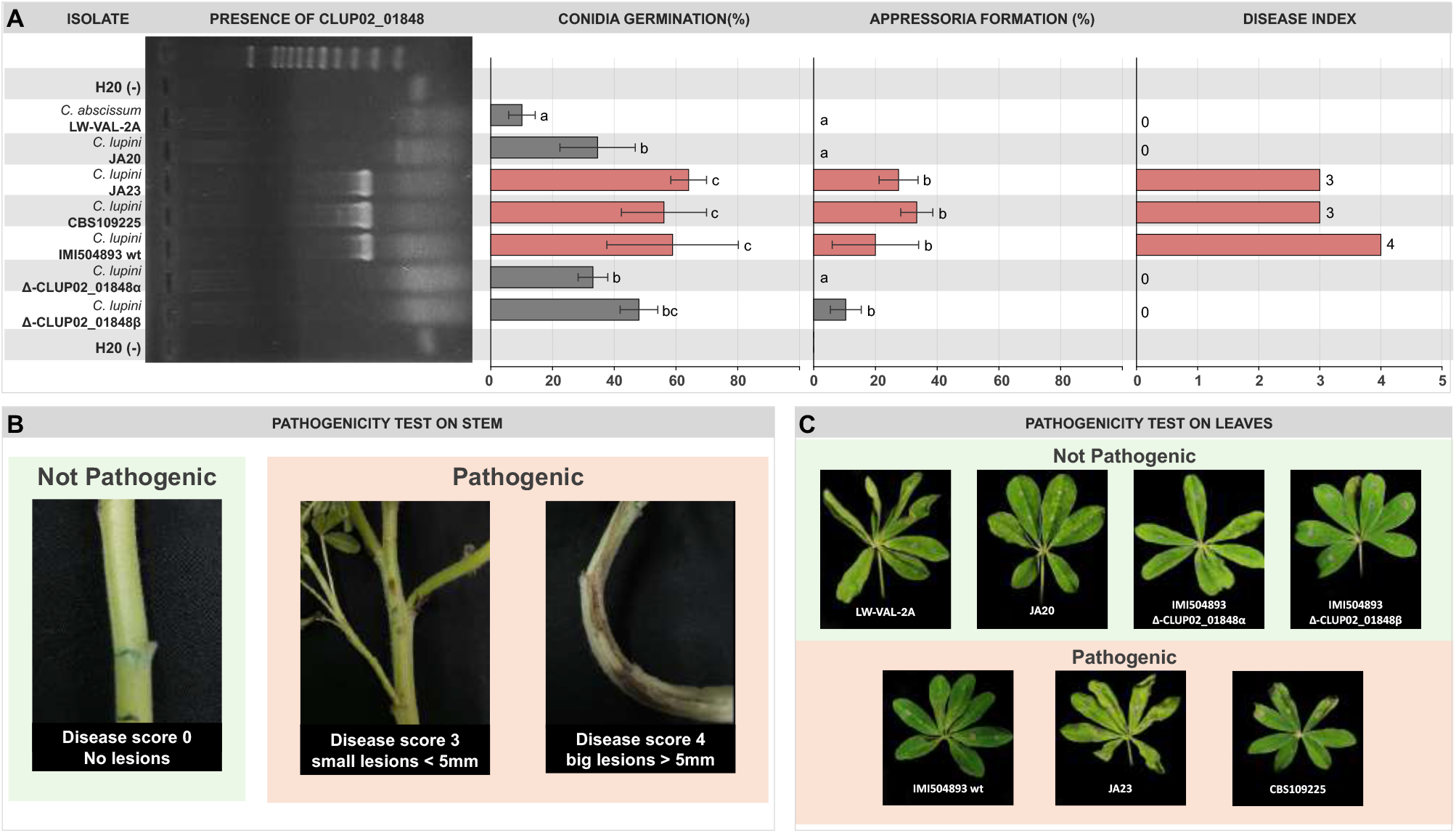
The PKS-NRPS biosynthetic gene cluster is required for pathogenicity. **A**, Molecular validation of PKS-NRPS backbone gene disruption and quantification of infection-related phenotypes. Left, PCR amplification of CLUP02_01848 showing the presence of the PKS-NRPS backbone gene in pathogenic *C. lupini* isolates and its absence in independent deletion mutants (Δ-CLUP02_01848α and Δ-CLUP02_01848β), as well as in non-pathogenic strains. Right, quantification of conidial germination, appressorium formation and disease index following inoculation of *Lupinus albus*. Bar plots show mean values for each isolate and mutant. Different letters indicate statistically significant differences among groups. Disease severity was scored ten days after inoculation using an ordinal disease index ranging from 0 (no lesions) to 4 (large lesions >5 mm). **B**, Pathogenicity assays on *Lupinus albus* stems inoculated with wild-type *C. lupini*, PKS-NRPS deletion mutants and non-pathogenic controls. Representative symptoms illustrate the progression from absence of lesions (disease score 0) to small lesions (<5 mm; disease score 3) and extensive necrotic lesions (>5 mm; disease score 4). Deletion mutants failed to induce disease symptoms comparable to those caused by the wild-type strain. **C**, Pathogenicity assays on *Lupinus albus* leaves following inoculation with wild-type *C. lupini*, PKS-NRPS deletion mutants and non-pathogenic controls. Representative leaf symptoms show that disruption of the PKS-NRPS backbone gene compromises pathogenicity, whereas wild-type pathogenic isolates induce characteristic necrotic lesions accompanied by abundant mycelial growth on the plant tissue surface.

Microscopic analyses revealed that the loss of pathogenicity was associated with defects in early infection-related development. Although conidial germination was observed, mutant strains displayed reduced germination rates and frequently produced abnormal, swollen germ tubes. Importantly, appressorium formation was severely impaired or absent, and when present, structures were weakly melanized compared with the fully melanized appressoria observed in the wild type. These defects indicate a failure to establish functional infection structures required for host penetration.

Consistent with these observations, both deletion mutants phenocopied the non-pathogenic *C. lupini* isolate JA20 and the non-pathogenic species *C. abscissum*, which also failed to form appressoria and did not cause disease. Together, these results demonstrate that the PKS-NRPS biosynthetic gene cluster is required for pathogenicity in *C. lupini*, establishing a direct functional link between this cluster and host infection.

### Population genomics reveals lineage-specific retention of derelict *Starship*s-associated genes

To assess the distribution of derelict *Starships* across the species, we analyzed genome variation in a global collection of *C. lupini* isolates (Fig. 1C & 7A). Population genomic analyses revealed a strongly structured population subdivided into four distinct lineages, with no evidence of recombination among them, consistent with a predominantly clonal population structure (Fig. 7A).

**Fig. 7.**
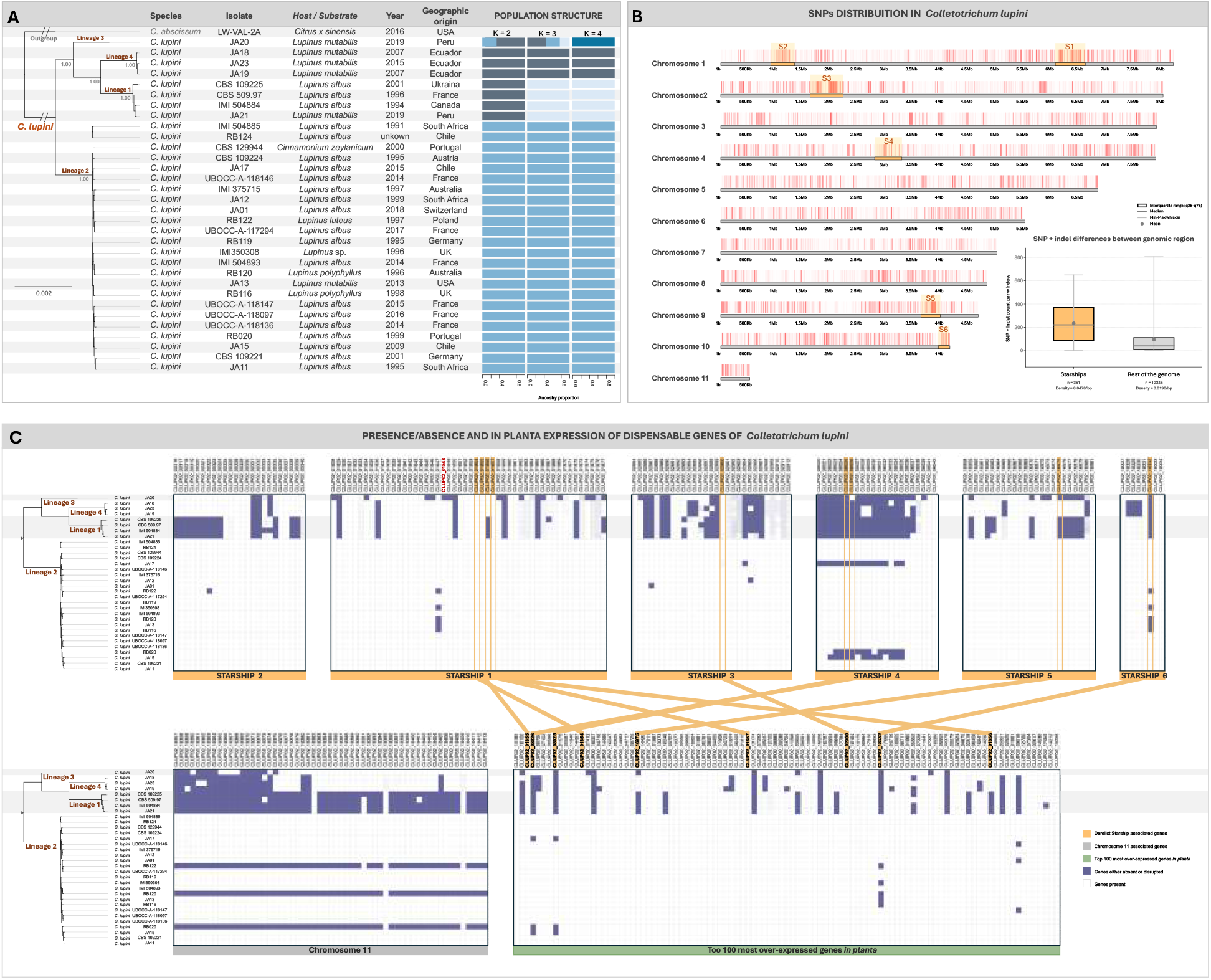
Lineage-specific variation of derelict *Starship*-associated genes in *C. lupini*. **A**, Phylogenomic relationships and ancestry composition of *C. lupini* isolates inferred from genome-wide sequence variation. Left, phylogenomic tree showing the subdivision of isolates into four major lineages. Right, population structure inferred for K = 2-4 genetic clusters, revealing lineage differentiation among isolates. Isolates are annotated by host or substrate of origin, year of isolation and geographic origin. *C. abscissum* was included as outgroup. **B**, Genome-wide distribution of SNPs and indels across *C. lupini* isolates. Variant density across the 11 chromosomes is shown together with the positions of derelict Starships (S1-S6). Boxplot summary statistics show that Starship regions accumulate substantially higher levels of sequence variation compared with the remainder of the genome. Boxes indicate interquartile ranges (q25-q75), centre lines indicate medians, whiskers indicate minimum and maximum values, and crosses indicate means. Variant density values for Starship regions and the rest of the genome are indicated. **C**, Presence-absence variation of genes associated with derelict *Starships* across *C. lupini* lineages. Heatmaps show the distribution of dispensable genes located within Starship regions and chromosome 11 across isolates. Purple and white cells indicate gene presence and absence or disruption, respectively. Several *Starship*-associated genes are absent or disrupted in the non-pathogenic isolate JA20. The lower panel shows the expression ranking of genes during plant infection, highlighting an enrichment of derelict Starship-associated genes among the top 100 most highly induced transcripts in planta. Orange lines connect Starship-associated genes with highly induced infection-related transcripts.

Variant analyses revealed that Starship regions exhibit markedly elevated levels of SNPs and indels compared with the rest of the genome, suggesting accelerated sequence evolution within these elements (Fig. 7B). Despite this limited genetic exchange, substantial variation was observed in the presence and absence of genes located within derelict *Starships*. In particular, genes within the *Starship 1* region, including those belonging to the PKS-NRPS biosynthetic gene cluster, were consistently retained in most pathogenic isolates but absent in the non-pathogenic isolate JA20. This pattern is consistent with a functional role for this region in host infection.

More broadly, derelict *Starships* showed lineage-specific patterns of retention and loss. Several regions were largely conserved within lineage II, the globally distributed pandemic lineage, but were partially or entirely absent in other lineages. Conversely, independent loss events were observed within lineage II, indicating ongoing turnover of these regions.

Consistent with their functional relevance, many genes located within derelict *Starships* are overrepresented among the most highly expressed infection-induced genes (Fig. 7C). In addition, we observed presence-absence polymorphism of an entire chromosome (chromosome 11) across the population, suggesting that chromosome 11 represents a dispensable chromosome and highlighting the remarkable genome plasticity of *C. lupini*. Together, these results indicate that *Starships* are subject to lineage-specific retention and loss and are enriched in infection-responsive genes. The conservation of the PKS-NRPS biosynthetic gene cluster across pathogenic isolates is consistent with a single acquisition event preceding lineage diversification. More broadly, these patterns support a model in which *Starship* regions contribute to standing genomic variation, with differential retention shaping lineage-specific genome composition.

## DISCUSSION

Our results identify a direct functional link between horizontally acquired genetic material and pathogenicity in a fungal plant pathogen. We show that a biosynthetic gene cluster embedded within a derelict *Starship* is required for host infection in *Colletotrichum lupini*, and that disruption of this cluster abolishes pathogenicity. By integrating comparative genomics, population genomics, transcriptomics, and functional genetics, we demonstrate that this region is transcriptionally active during infection, exhibits atypical evolutionary signatures, and is differentially retained across lineages. Together, these findings suggest that derelict *Starships* can act as reservoirs of adaptive genetic variation and harbour key traits associated with host specialization. Considering the expansion of derelict *Starships* in *C. lupini*, the strong induction of several genes located within these regions during plant infection, and the temporal overlap between major evolutionary events, it is intriguing to speculate that *Starships* may have fuelled both the acquisition and genomic compartmentalization of these regions, thereby promoting the emergence and maintenance of adaptive pathogenicity-associated traits.

Placing these observations in an evolutionary context suggests that the emergence of pathogenicity may have coincided with the arrival of a new host species within a natural ecological setting. Divergence time estimates indicate that *C. lupini* split from its closest known relative approximately 1.87 million years ago, closely overlapping with the emergence of *Lupinus* species in western South America^32,33^. This temporal concordance is consistent with an early host-association event within this geographic region. Notably, subsequent diversification within *C. lupini* overlaps with the radiation of its lupin hosts^32,33^, suggesting that the evolutionary trajectory of this interaction likely involved prolonged host adaptation and co-diversification rather than a single discrete host jump. Such dynamics are consistent with evolutionary models in which plant pathogens undergo host specialization and genetic isolation following initial host association^5–9^.

Interestingly, *C. lupini* is capable of infecting a broad range of lupin species, including *Lupinus albus*, the cultivated white lupin, which belongs to an Old World lineage that diverged prior to the diversification and radiation of New World lupin species^32,33^. This broad host compatibility suggests that *C. lupini* may have acquired the ability to overcome defense mechanisms conserved across the *Lupinus* genus, thereby facilitating host colonization across phylogenetically divergent lupin species.

Within this framework, the PKS-NRPS biosynthetic gene cluster identified here likely represents a key innovation associated with host colonization. Its conservation across pathogenic isolates, absence in closely related non-pathogenic species, and requirement for infection support a model in which this cluster was acquired prior to lineage diversification and subsequently maintained under strong selective constraint. Although homologues of the PKS-NRPS backbone gene are distributed across diverse fungal lineages, the phylogenetic incongruence observed here, together with the absence of the cluster from closely related taxa, is consistent with horizontal acquisition of this locus. More broadly, the complex evolutionary history of these genes suggests that gene mobility has contributed to their distribution across distant fungal clades^13–15,17,34–36^.

At the genomic level, our analyses reveal a compartmentalized organization in *C. lupini*, consisting of a conserved core genome and a dynamic compartment enriched in transposable elements and lineage-specific genes, and characterized by several genes that are highly induced during interaction with the plant^11,13^. This organization reflects a continuum of genomic adaptation, in which rapidly evolving regions contribute disproportionately to host interaction and adaptation. Our results further indicate that these highly dynamic regions are associated with derelict *Starships*, suggesting that *Starships* have contributed to shaping genome architecture and adaptive potential. Similar elements have been shown to mediate large-scale gene transfer and genome restructuring in fungi^22–24^. Importantly, all identified *Starships* lack intact recombinase catalytic domains and show signatures of structural decay, indicating that they represent degenerated remnants of ancestrally active elements rather than currently mobile entities. In this context, the current genome architecture likely reflects the legacy of past episodes of genome-scale mobility followed by long-term stabilization and degeneration.

A key feature of these derelict *Starships* is their enrichment in genes with atypical evolutionary signatures and strong transcriptional induction during infection. Horizontal gene transfer has long been recognized as a major source of innovation in fungal genomes, particularly for genes involved in host interaction and secondary metabolism^13–15,17,34–36^. However, our analyses indicate that, within *C. lupini*, robust evidence for horizontal acquisition is restricted to the backbone gene of the PKS-NRPS biosynthetic cluster. Other genes within these regions may reflect more complex evolutionary processes, including differential retention, sequence divergence or incomplete lineage sorting. This distinction highlights the importance of combining phylogenetic and functional evidence when inferring horizontal transfer.

Beyond their role in gene acquisition, *Starships* also contribute to genomic variation within the species. Population genomic analyses revealed lineage-specific patterns of retention and loss, with the globally distributed pandemic lineage retaining a more complete complement of these regions. In contrast, partial or complete loss of these regions in other lineages, including the non-pathogenic isolate JA20, indicates ongoing turnover. These patterns are consistent with a model in which derelict *Starships* represent a source of standing genomic variation, with differential retention contributing to lineage-specific genomic composition. Based on the evolutionary analyses, it is likely that JA20 represents secondary loss of pathogenicity rather than a retention of an ancestral non-pathogenic state. More broadly, our findings highlight the potential role of *Starships* in facilitating genome evolution in filamentous fungi. While transposable elements have long been recognized as drivers of genome plasticity and host-pathogen coevolution^20^, the contribution of giant cargo-carrying elements to the assembly and dissemination of complex gene repertoires is only beginning to be understood^22–24^. Our results provide functional evidence linking such a region to a key pathogenicity determinant, supporting the view that genome-scale mobility can contribute to the emergence of adaptive traits.

Several questions remain. The mechanisms underlying the mobilization and transfer of these elements are not fully resolved, and the extent to which they move within or between species remains unclear. In addition, although we demonstrate a central role for a single biosynthetic gene cluster, the functional contributions of other genes located within derelict *Starships* remain to be determined. Addressing these questions will be essential for understanding how genome-scale processes contribute to the emergence and diversification of fungal pathogens^1–4^.

Together, our results support a model in which *Starships* act as catalysts of genome innovation during the evolution. Although these elements are now degenerated and immobile, their legacy persists in the form of structurally and functionally distinct genomic compartments that harbour key adaptive traits. We propose that early genome-scale mobility facilitated the acquisition and integration of complex functions required for host association, whereas subsequent evolutionary trajectories were shaped by differential retention, loss and functional refinement of these regions. In this view, derelict *Starships* represent enduring genomic imprints of past mobility events that continue to structure the adaptive landscape of fungal pathogens.

## MATERIALS AND METHODS

### Fungal isolates and plant material

A collection of 65 *Colletotrichum* strains was assembled to represent the phylogenomic diversity of the genus, with denser sampling of *C. lupini* isolates spanning its geographic and lineage diversity (Supplementary File S10). Plant infection assays were performed using *Lupinus albus* cultivars under controlled growth conditions.

### Pathogenicity assays

Pathogenicity was assessed using seedling, detached leaf and stem inoculation assays. Conidial suspensions were prepared from cultures grown on potato dextrose agar and adjusted to standardized concentrations prior to inoculation. Disease development was monitored over time, and infection phenotypes were evaluated based on lesion formation and disease severity. Detailed experimental conditions are provided in Supplementary File S10.

### Genome sequencing and assembly

Genomic DNA was extracted from fungal cultures and sequenced using Illumina paired-end technology. Reads were quality filtered and assembled *de novo*, and genome completeness was assessed using standard benchmarking approaches. Gene prediction was performed using *ab initio* and evidence-based methods. Detailed pipelines and parameters are described in Supplementary File S10.

### Comparative genomics and phylogenomics

Orthology inference and phylogenomic reconstruction were performed using genome-wide protein datasets. Single-copy orthologs were aligned and used to infer species relationships and divergence times. Gene presence-absence and copy-number variation were analyzed to identify lineage-specific gene content. Detailed methods are provided in Supplementary File S10.

### Identification of *Starships*

Dynamic genomic regions were identified through comparative genomics and synteny analyses. Candidate *Starships* were annotated using a combination of automated detection and manual curation based on structural and sequence features, including the presence of recombinase-related genes. Transposable elements were annotated using established pipelines. Full methodological details are provided in Supplementary File S10.

### Transcriptomic analysis

Publicly available RNA-seq data were analyzed to assess gene expression during plant infection. Reads were aligned to the reference genome and differential expression analyses were performed to compare in planta and in vitro conditions. Detailed analytical steps and statistical thresholds are described in Supplementary File S10.

### Population genomics

Genome variation across *C. lupini* isolates was analyzed using read mapping and variant calling approaches. Population structure was inferred using multivariate and clustering methods. Gene presence-absence and mutation matrices were generated to assess variation in *Starships*. Detailed procedures are described in Supplementary File S10.

### Gene disruption and phenotypic analysis

The PKS-NRPS backbone gene was disrupted using Agrobacterium-mediated transformation. Independent deletion mutants were obtained and validated, and their phenotypes were assessed using pathogenicity assays and microscopy. Detailed protocols are provided in Supplementary File S10.

## ACKNOWLEDGEMENTS

This research and the APC were partially supported by: (1) the PROGRAILIVE project (grant number RBRE160116CR0530019), funded by the regions of Bretagne and Pays de la Loire (France) and the European Union (FEADER); (2) by the AsCoLuP project (*Ascochyta Colletotrichum Lupin Pois chiche*, CASDAR n°19AIP5913); (3) the OXIGEN project (grant number PID2024-161771NB-I00, AEI/10.13039/501100011033/FEDER, EU), funded by the Ministry of Science, Innovation and Universities (Spain).

We thank the curators and staff of the UBOCC (Université de Bretagne Occidentale Culture Collection), and in particular Amélie Weill and Patrice Nodet, as well as the CABI and CBS culture collections, for kindly providing the strains used in this study. We also thank Joris Alkemade, Monika Messmer, and Pierre Hohmann (Department of Crop Sciences, Research Institute of Organic Agriculture (FiBL), Frick, Switzerland), and Liliana Cano (Florida Research Center for Agricultural Sustainability, Vero Beach, FL, United States), for providing fungal isolates. Finally, we thank Antonio Prodi, Samantha Zoë Thomissen, and the entire Plant Pathology group at University of Bologna for technical and logistical support.

## EXTENDED DATA

**Supplementary File S1. List of fungal strains and genome assemblies used in this study**. This supplementary file contains detailed information on the fungal strains and genome assemblies analyzed in this study, including species identification, species complex assignment, host or substrate of origin, geographic origin, year of isolation, sequencing technology, genome accession numbers, assembly statistics, and BUSCO completeness metrics.

**Supplementary File S2. Functional annotation of selected orthogroups identified in this study**. This supplementary file contains the annotation of selected orthogroups identified through comparative genomic analyses. Information includes orthogroup identifiers, predicted gene functions, conserved protein domains, functional categories, and associated annotations derived from sequence similarity and bioinformatic analyses.

**Supplementary File S3. Functional annotation of predicted genes across *Colletotrichum lupini* IMI504893 genome**. This supplementary file contains the functional annotation of predicted genes identified in the genomes analyzed in this study. The dataset includes gene identifiers, predicted protein functions, conserved domains, Gene Ontology (GO) terms, enzyme classifications, and annotations obtained from homology-based and domain-based bioinformatic analyses.

**Supplementary File S4. Transposable element-mediated gene fragmentation in *Colletotrichum lupini***. This image file illustrates a case of gene fragmentation associated with the insertion of a transposable element (TE) in *Colletotrichum lupini*. The figure highlights the genomic organization surrounding the disrupted locus, including intact and fragmented genes, TE annotations, and neighboring coding sequences, providing evidence for TE-mediated gene disruption.

**Supplementary File S5. Differential gene expression analysis of *Colletotrichum lupini* during infection versus liquid culture**

This supplementary file contains the DESeq2 results from the comparative transcriptomic analysis of *Colletotrichum lupini* grown under infection conditions versus liquid-culture controls. The dataset includes normalized expression values, log_2_ fold changes, adjusted p-values, and statistical significance for genes identified as differentially expressed during host infection. Detailed materials and methods for RNA-seq processing, read mapping, differential expression analysis, and data visualization are provided in the Supplementary Materials and Methods document.

**Supplementary File S6. Comparative structure and synteny of a horizontally transferred biosynthetic gene cluster across fungal genomes** This supplementary file illustrates the genomic organization and synteny of a putatively horizontally transferred PKS/NRPS biosynthetic gene cluster (BGC) identified in *Colletotrichum lupini* and distantly related fungal species. The figure compares gene content, orientation, and conservation across multiple genomes, highlighting the structural conservation and evolutionary relationships of the BGC region.

**Supplementary File S7. Presence/absence patterns and protein conservation scores of selected genes across *Colletotrichum* isolates** This supplementary file reports the distribution and conservation status of selected genes, identified by locus tag, across all analyzed isolates. For each isolate, a value of 1 indicates the presence of a fully conserved gene, whereas 0 indicates either gene absence or the presence of a predicted deleterious mutation affecting the encoded protein. Intermediate represent the protein conservation score, reflecting the degree of sequence conservation relative to the reference sequence, with higher values indicating greater conservation. The dataset was generated by integrating gene presence/absence analyses with predicted functional impacts of sequence variants.

**Supplementary File S8. Population structure and SNP-based genomic analyses of the analyzed *Colletotrichum lupini* isolates**. (A) Distribution of Cross-validation Error (CV error) values calculated to identify the optimal number of genetic clusters (K) for the population structure analysis. The selected K corresponds to the model with the best-supported population partitioning. (B) Principal Component Analysis (PCA) based on genome-wide single nucleotide polymorphisms (SNPs), illustrating the genetic relationships and clustering patterns among isolates. (C) SNP-based network analysis showing the genetic connectivity and relatedness among isolates inferred from pairwise genomic variation. Distinct genetic lineages identified through population genomic analyses are highlighted.

**Supplementary File S9. Pathogenicity phenotypes caused by *Colletotrichum* isolates on lupin tissues**. Representative disease symptoms observed on lupin tissues following inoculation with different *Colletotrichum* isolates and mutant strains. The figure illustrates variation in pathogenicity and disease severity among isolates, including wild-type and gene-disruption mutants, based on lesion development and symptom progression on infected plant tissues.

**Supplementary File S10. Supplementary materials and methods**. This supplementary file provides detailed descriptions of the experimental procedures, bioinformatic pipelines, comparative genomic analyses, transcriptomic analyses, population genomic approaches, and functional validation experiments performed in this study. Methods related to pathogenicity assays, genome sequencing and assembly, transposable element annotation, phylogenomics, RNA-seq differential expression analyses, SNP analyses, and targeted gene disruption are described together with the software, parameters, and statistical approaches used throughout the study.

