## Supplementary File S4. for "Adaptive genomic compartments shaped by giant mobile elements underpin the ancient emergence of fungal pathogenicity": Supplementary File S4. Transposable element-mediated gene fragmentation in Colletotrichum lupini.pdf

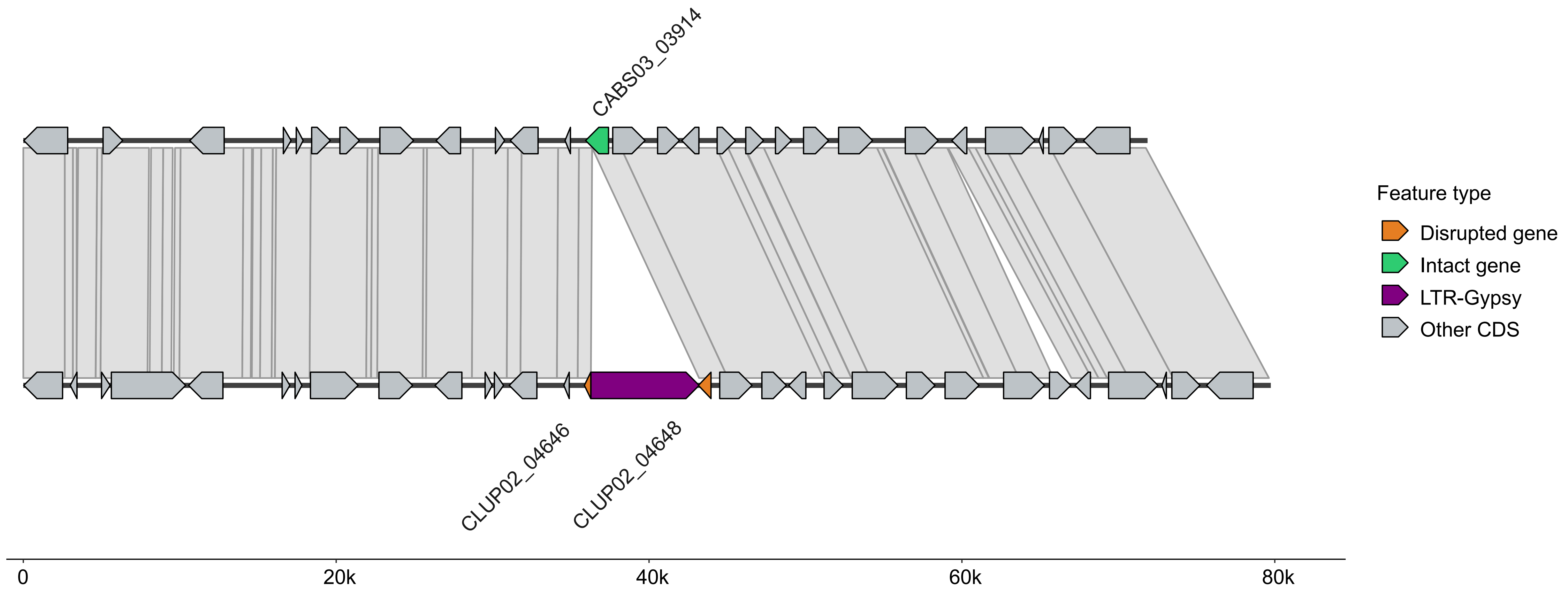


**Supplementary File S4. Transposable element-mediated gene fragmentation in *Colletotrichum lupini*.** This image file illustrates a case of gene fragmentation associated with the insertion of a transposable element (TE) in *Colletotrichum lupini*. The figure highlights the genomic organization surrounding the disrupted locus, including intact and fragmented genes, TE annotations, and neighboring coding sequences, providing evidence for TE-mediated gene disruption.
