## Supplementary File S6. for "Adaptive genomic compartments shaped by giant mobile elements underpin the ancient emergence of fungal pathogenicity": Supplementary File S6. Comparative structure and synteny of a horizontally transferred biosynthetic gene cluster across fungal genomes .pdf

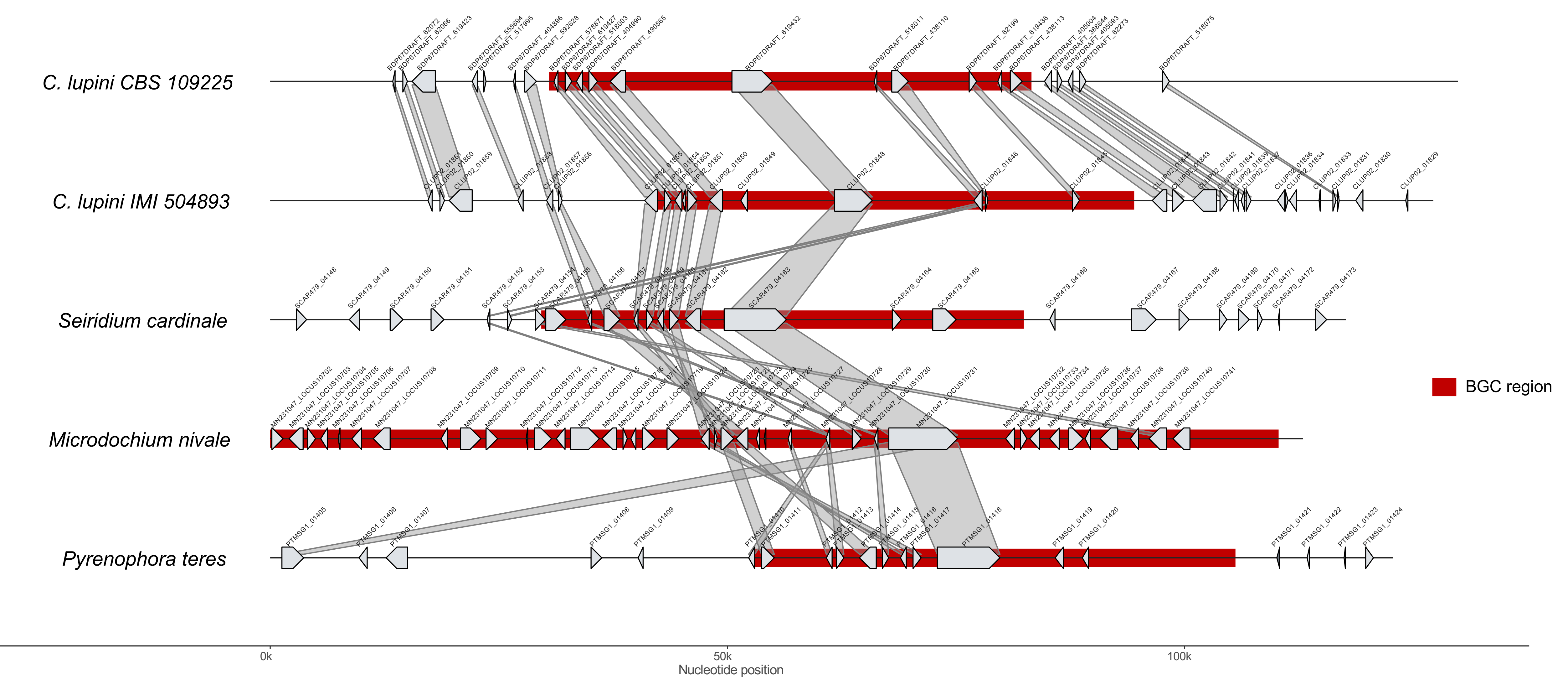


**Supplementary File S6. Comparative structure and synteny of a horizontally transferred biosynthetic gene cluster across fungal genomes**

This supplementary file illustrates the genomic organization and synteny of a putatively horizontally transferred PKS/NRPS biosynthetic gene cluster (BGC) identified in *Colletotrichum lupini* and distantly related fungal species. The figure compares gene content, orientation, and conservation across multiple genomes, highlighting the structural conservation and evolutionary relationships of the BGC region.
