## Supplementary File S8. for "Adaptive genomic compartments shaped by giant mobile elements underpin the ancient emergence of fungal pathogenicity": Supplementary File S8. Population structure and SNP-based genomic analyses of the analyzed Colletotrichum lupini isolates.pdf

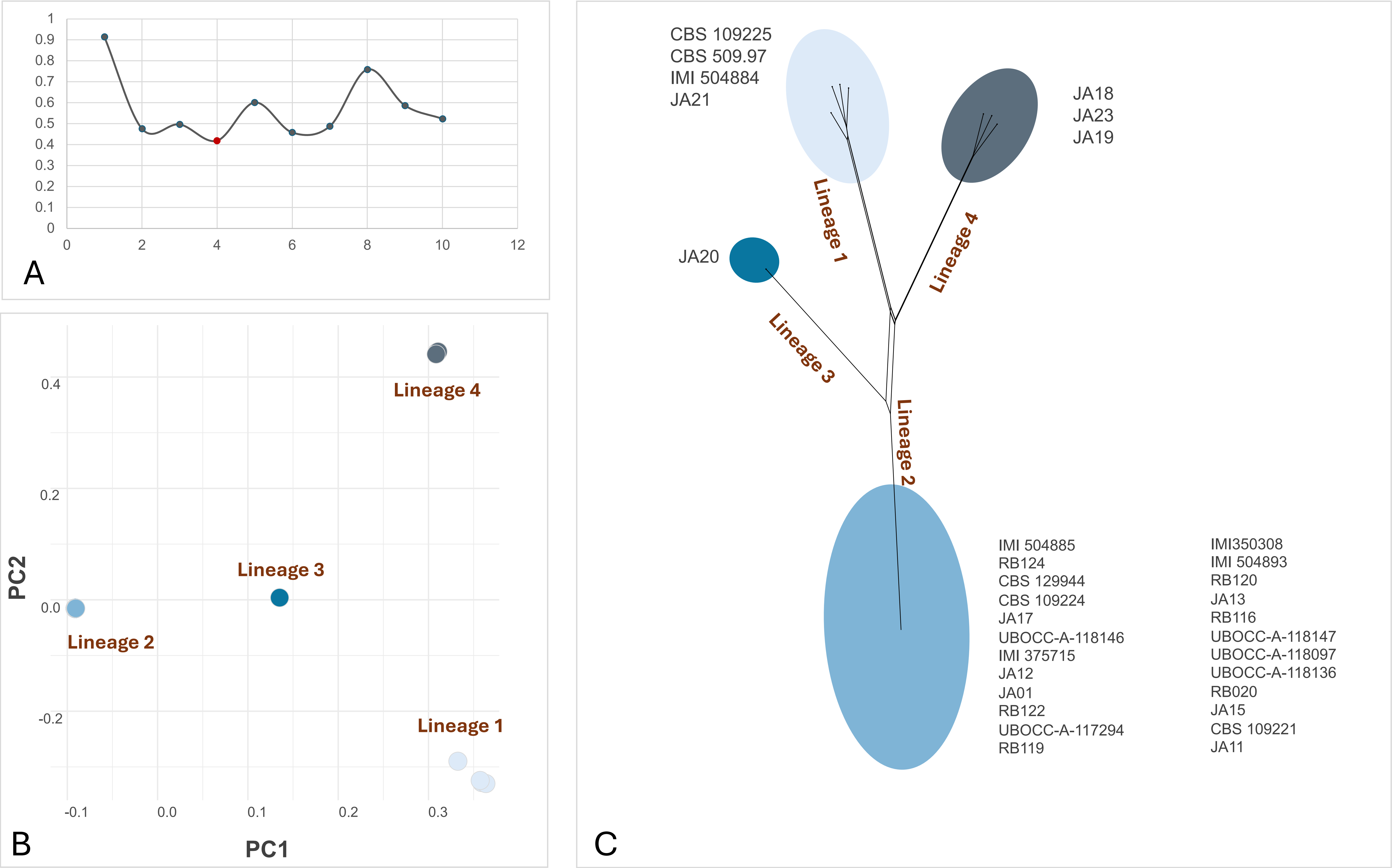


**Supplementary File 8. Population structure and SNP-based genomic analyses of the analyzed *Colletotrichum lupini* isolates**

(A) Distribution of Cross-validation Error (CV error) values calculated to identify the optimal number of genetic clusters (K) for the population structure analysis. The selected K corresponds to the model with the best-supported population partitioning.

(B) Principal Component Analysis (PCA) based on genome-wide single nucleotide polymorphisms (SNPs), illustrating the genetic relationships and clustering patterns among isolates.

(C) SNP-based network analysis showing the genetic connectivity and relatedness among isolates inferred from pairwise genomic variation. Distinct genetic lineages identified through population genomic analyses are highlighted.
