## Supplementary File S9. for "Adaptive genomic compartments shaped by giant mobile elements underpin the ancient emergence of fungal pathogenicity": Supplementary File S9. Pathogenicity phenotypes caused by Colletotrichum isolates on lupin tissues.pdf

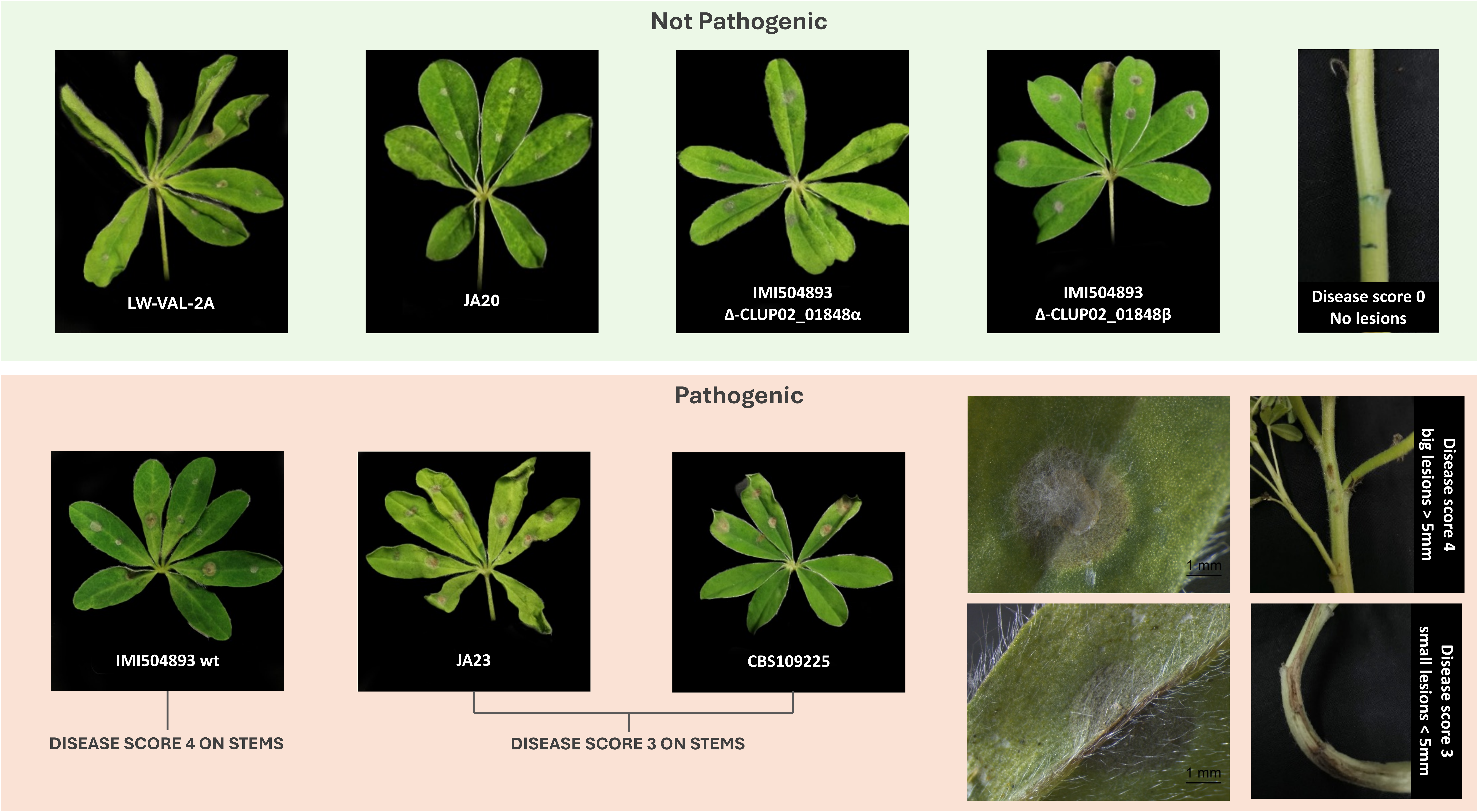


**Supplementary File S9. Pathogenicity phenotypes caused by Colletotrichum isolates on lupin tissues**. Representative disease symptoms observed on lupin tissues following inoculation with different *Colletotrichum* isolates and mutant strains. The figure illustrates variation in pathogenicity and disease severity among isolates, including wild-type and gene-disruption mutants, based on lesion development and symptom progression on infected plant tissues.
