## Supplementary File S10. for "Adaptive genomic compartments shaped by giant mobile elements underpin the ancient emergence of fungal pathogenicity": Supplementary File S10. Supplementary materials and methods..pdf

### SUPPLEMENTARY MATERIAL AND METHODS

#### Fungal isolates and plant material

To investigate the evolutionary origin of *Colletotrichum* pathogenicity on lupin, we assembled a collection of 65 strains representing the phylogenomic diversity of the genus, with denser sampling of the focal lineage. The dataset included multiple *C. lupini* isolates spanning the species' geographic and phylogenomic diversity[1,2], representatives of all recognized species within Clade 1 of the *C. acutatum* species complex[3,4], and additional polyphagous and more distantly related *Colletotrichum* species[5,6]. Strain selection prioritized the availability of high-quality genome assemblies to support robust phylogenomic reconstruction and comparative genomic analyses. Details of all strains are provided in Supplementary File 1.

#### Pathogenicity and virulence assays

Seed-inoculation pathogenicity assays

All isolates (Fig. 1 and Supplementary File 1) were subcultured on potato dextrose agar (PDA; Difco, USA) for 10–14 days at 25 ± 1 °C before pathogenicity assays. Conidia were harvested by flooding plates with sterile distilled water, gently scraping the colony surface, and filtering the suspension through sterile gauze to remove mycelial fragments. Spore concentrations were adjusted to $1 \times 10^6$ conidia $mL^{-1}$ using a haemocytometer.

Seeds of *Lupinus albus* 'Energy' were surface-sterilized in 5% sodium hypochlorite for 3 min, rinsed with sterile distilled water, and germinated in sterile Petri dishes containing autoclaved sand at field capacity. Germinated seeds (2–3 days old) were immersed in the conidial suspension for 2 min before transfer to sterile substrate. Control seeds were treated with sterile water. Seedlings were maintained at 25 ± 2 °C under a 16 h light/8 h dark photoperiod. Disease severity was evaluated 21 days post-inoculation using ten biological replicates per isolate. Pathogenicity was determined based on anthracnose symptoms and/or seedling mortality, and Koch's postulates were confirmed by re-isolation of the fungus from symptomatic tissues.

Pathogenicity test on lupin leaves

Fully expanded detached leaves of *Lupinus albus* 'Misak' were inoculated on the adaxial surface of the leaflet blade with 10 µL droplets containing $1 \times 10^6$ conidia $mL^{-1}$ supplemented with 1% gelatin to promote adhesion to the leaf surface. Conidial suspensions were prepared from 10-day-old cultures grown on potato dextrose agar (PDA) by flooding plates with sterile distilled water, gently scraping the colony surface, filtering the suspension through cheesecloth, and adjusting the concentration after counting with a haemocytometer. One or two droplets were deposited per leaflet on four leaves per isolate. Inoculated leaves were incubated in a moist chamber at 25 °C under a 12 h photoperiod. Conidial germination and appressorium formation were assessed 24 h after inoculation. Leaf imprints were obtained by applying transparent nail polish to the inoculation sites[7#]. After drying, the nail-polish film was removed, stained with cotton blue, and examined under a light microscope (Leitz Laborlux S) at 400× magnification. For each isolate, 100 conidia were examined in each of eight droplets. Conidial germination was calculated as the percentage of germinated conidia relative to the total number observed, and appressorium formation was expressed as the proportion of germinated conidia that developed appressoria. Disease symptoms were monitored on inoculated leaves for up to 8 days after inoculation. Statistical analyses were performed using Tukey's test ($\alpha = 0.05$) implemented in Statistica 8.0 (StatSoft, USA).

<u>Pathogenicity test on lupin stems</u>

Inoculation conditions were identical to those used for leaf inoculation. For each isolate, eight to ten inoculation spots were selected on stems of *Lupinus albus* 'Misak' at bloom stage, and at each site a 10 µL droplet containing $1 \times 10^6$ conidia $mL^{-1}$ and 1% gelatin was deposited with a micropipette. Stems were not wounded, following Thomas et al.2008[8]. Plants were incubated for 24 h in a wet chamber at 25 °C under a 12 h photoperiod and then maintained under similar temperature and photoperiod conditions at ambient humidity. Symptoms were assessed 10 days after inoculation. The most severe stem lesion on each plant was scored using a 0–5 disease severity scale, where 0 = no lesion present; 1 = distinct stem bending or pinpoint lesion less than 1 mm in diameter; 2 = small lesion less than 5 mm in diameter, no sporulation; 3 = small lesion less than 5 mm in diameter with sporulation; 4 = large lesion girdling more than half the circumference of the stem with abundant sporulation; and 5 = large lesion severing the stem[8,9].

**<u>DNA extraction, genome sequencing, assembly and gene prediction</u>**

All isolates were grown on Potato Dextrose Agar (PDA) for 7 days before DNA extraction. Mycelium was scraped from colony surfaces and genomic DNA was extracted using a modified CTAB protocol. Samples were homogenized in 700 µL of 3% CTAB using a Motor-Driven Tissue Grinder G50 (Coyote Bioscience, Beijing, China) and incubated at 65 °C for 30 min. DNA was purified by chloroform:isoamyl alcohol extraction (24:1) and precipitated with ice-cold isopropanol at −20 °C overnight. The DNA pellet was washed with 70% ethanol, resuspended in 50 µL nuclease-free water, quantified using a NanoDrop ND-1000 spectrophotometer (Thermo Scientific, USA), and stored at −20 °C until sequencing.

Whole-genome sequencing was performed on an Illumina NovaSeq 6000 platform using 150 bp paired-end reads. Read quality was assessed with FastQC v0.12.1 and adapters and low-quality bases were removed using Trimmomatic v0.33[10]. Reads were merged with FLASH v1.2.11[11] and assembled using SPAdes v3.15.1[12]. Low-coverage scaffolds were removed as potential contaminants. Assembly quality and completeness were evaluated using QUAST v5.0.2[13] and BUSCO v3.1[14]. Repetitive sequences were identified using RepeatModeler v2.0.6[15] and RepeatMasker v4.1.5[16]. Gene prediction was performed with AUGUSTUS v3.5.0[17] using a species-specific model trained with publicly available *C. lupini* RNA-seq data (SRX2782478[18]) assembled with rnaSPAdes v3.15.1[19]. Protein-coding sequences were extracted from the GFF3 output using *getAnnoFasta.pl* (available in AUGUSTUS v3.5.0[17]).

Identification and characterization of genes and gene families

For identification and characterization of specific gene families, sequences were annotated using the JGI annotation pipeline[20]. Secreted proteins were predicted using DeepTMHMM v1.0[21]. Protein domains and functional annotations were assigned using Pfam[22] and InterPro terms[23]. Carbohydrate-active enzymes (CAZymes) were identified using the CAZy annotation pipeline[24], while peptidases were annotated using the MEROPS database[25].

Genes encoding backbone enzymes of secondary metabolite biosynthetic clusters, including nonribosomal peptide synthetetases, polyketide synthases, DMATS-family aromatic prenyltransferases, and terpene synthases, were identified using BLASTp and InterProScan domain annotations; biosynthetic gene clusters were identified and annotated with Antismash 8.0.4[26].

Transcription factors were identified by BLASTp[27] searches against the NCBI nr database and the Aspergillus Genome Database (AspGD) using a threshold of 1e−10[28]. Functional domains were verified using the Conserved Domain Database (CDD) and SMART[29].

Phylogenomic reconstruction and divergence time estimation

Phylogenomic analyses were performed following the pipeline described as follows.[30] Predicted proteomes were clustered into orthologous groups using OrthoFinder v0.4[31]. Single-copy Orthogroups were retrieved and protein sequences aligned using MAFFT v7.525[32], refined with Gblock v0.91[33] and concatenated using AMAS v0.98[34]. Phylogenomic trees were constructed from the concatenated alignment using RAxML v8.2.11[35], FastTree v2.1.11[36] and MrBayes v3.2.6[37].

A calibrated timetree was inferred using the RelTime method implemented in MEGA X v10.1.7[38], using the crown of *Colletotrichum* at 68.76 MYA (103.85–45.53 MYA)[39] as calibration point.

#### Transposable element and repetitive sequence annotation

##### Identification of repetitive elements

Transposable elements (TEs) and repetitive sequences were annotated *de novo* in all assemblies using EarlGrey v6.2.0[40], which integrates RepeatModeler2[15], RepeatMasker[41], BLAST+[27], MAFFT[32], trimAl[42], LTRharvest[43], LTR_retriever[44], RepeatCraft[45], and Tandem Repeats Finder[46]. Each genome assembly was analysed using the Dfam eukaryotic partition as a reference library together with a de novo repeat library generated by RepeatModeler2. TEstrainer[47] iteratively refined consensus sequences through cycles of BLAST-based extraction, alignment with MAFFT v7.525[32], and trimming with trimAl[42]. Final annotations were defragmented using RepeatCraft[45], and repeat landscape analyses were enabled (-e yes). TEs were classified according to the unified system of Wicker et al.[48]. Repeat landscapes were visualized as CpG-corrected Kimura divergence plots to estimate the relative age and activity of TE families. The resulting TE annotations were used for TE density calculations and integrated with *Starship* element boundaries identified using the Starfish annotation workflow described below.

##### Annotation of giant cargo-mobilizing transposable elements

To characterize giant cargo-mobilizing transposable elements, including *Starships* and their associated tyrosine recombinase genes ("captains"), we used Starfish v1.1.0[49], a dedicated pipeline for identifying *Starship* and captain-like tyrosine recombinases (YR).

The candidate captains (YR genes) were identified de novo from the genomes using the Starfish gene finder module with MetaEuk[50] and HMMER[51] based on sequence similarity to the Pezizomycotina YR database provided by Starfish. The resulting YR list for *C. lupini* was inspected for membership in species-specific genes or expanded gene families identified by orthogroup analysis.

Candidate *Starship* elements were initially identified based on large genomic gaps detected in whole-genome alignments generated with SyMAP v5.6.0[52] and MAUVE v2.3.1[53] (further details in *Synteny Analysis* section), together with the presence of YR genes. Resulting elements were evaluated with Starfish-detected *Starship* candidates that lacked flanking synteny against orthologous sites without *Starships* through manual inspection using the command pairViz[49,54], and by intersecting the architecture of Starfish-identified elements with the previously defined species-specific genes and regions.

#### Comparative genomics

##### Protein clustering

Orthology-based clustering of predicted proteomes was performed using OrthoFinder v2.5.2[31] across 24 *Colletotrichum* genomes, including the reference *C. lupini* isolates (CBS109225, RB173 and IMI04893, RB221) and representatives of 20 additional species, with *Verticillium dahliae* used as an outgroup. Orthogroup presence/absence and copy-number variation were analyzed to identify gene families specific to, expanded in, or contracted in the *C. lupini* reference genome.

##### Lineage-specific region (LSR) identification

To identify genomic regions unique to *C. lupini*, raw reads from 23 isolates (Supplementary File 1) were aligned to the IMI 504893 reference genome using Bowtie v1.3.1[55]. Mean coverage was calculated in 5-kb non-overlapping windows, and regions with average coverage <2× in all non-*C. lupini* isolates were flagged as missing and therefore considered candidate *C. lupini* lineage-specific regions (LSRs). Candidate regions were further validated by BLASTn searches against all assembled genomes to confirm the absence of homologous sequences in non-*C. lupini* genomes.

##### Synteny analysis

Whole-genome synteny was assessed between the *C. lupini* reference genome and two high-quality, near-chromosome-level assemblies (*C. abscissum* LW-VAL-2A and *C. paranaense* IMI 384185) using SyMAP v4.2.42[56]. Collinear chromosomes were subsequently aligned using Mauve v2.4.1[53].

#### Meta-analysis of *C. lupini* RNA-seq datasets and expression profiles

Raw RNA-seq reads from *C. lupini* strain IMI504893 also known as RB221 (GCF_023278565.1[18]) were quality-checked using FastQC v0.11.7[57] and summarized with MultiQC v1.28[58]. Low-quality bases and adapters were trimmed using Trimmomatic v0.27[10], applying end trimming, sliding-window quality filtering, and a minimum read length of 50 bp. Read quality was re-evaluated with MultiQC. Splice sites and exon coordinates were extracted from the NCBI GTF annotation to build a genome index for HISAT2 v2.2.1[59]. Strand-specific reads were aligned to the reference genome using HISAT2 in reverse-forward stranded mode, and resulting alignments were processed with SAMtools v1.21[60].

Gene-level read counts were generated using featureCounts v2.0.8[61] in paired-end, reverse-stranded mode, assigning reads to genes based on exon features and gene_id annotations. The final count matrix comprised 24 samples spanning liquid-culture and infection timepoints.

Differential expression analyses were conducted in R v4.3.2[62] using DESeq2 v1.42.1[63]. After filtering genes with zero counts across all samples, infection samples were compared collectively against liquid-culture controls using a design formula of ~ group. $\text{Log}_2$ fold changes were shrunk using apeglm v1.32.0 [64] to stabilize effect-size estimates. Genes with adjusted $p \leq 0.05$ and $|\text{shrunken } \log_2\text{FC}| \geq 1$ were considered significantly differentially expressed.

Volcano plots for the pooled Infection versus liquid-culture comparison were generated with ggplot2 v3.5.2[65]. In one representation, predefined custom gene bundles were overlaid on the transcriptome-wide background, whereas in another, genes located within curated derelict *Starship* loci were highlighted.

To investigate temporal dynamics, a second DESeq2 model was applied to compare individual infection timepoints (24, 48, 60, 72, and 84 hpi) against liquid-culture controls. Variance-stabilized expression values were obtained using the DESeq2 vst() function. Expression patterns of genes within curated derelict *Starship* with 50 upstream and 50 downstream flanking genes were visualized using ComplexHeatmap v2.18.0[66]. Statistical significance derived from the DESeq2 differential expression analyses (adjusted $p \leq 0.05$) was additionally annotated alongside the heatmaps as a binary indicator (significant vs non-significant).

#### **Population genomics**

##### Variant detection and filtering

Single-nucleotide polymorphisms (SNPs) and short insertions/deletions (indels) were identified by mapping sequencing reads to the *Colletotrichum lupini* IMI 504884 reference genome using BWA v0.7.17[67]. SAM files were converted to BAM format, sorted and indexed using SAMtools v1.17[68]. Variant calling was performed using GATK v4.4.0[69]. HaplotypeCaller, followed by CombineGVCFs and GenotypeGVCFs, with ploidy set to 1 and a maximum of two alternative alleles. Variants were filtered using hard-filter thresholds based on GATK quality metrics to remove potentially erroneous calls. Site-level filters included Fisher strand bias (FS > 60), mapping quality (MQ < 40), quality by depth (QD < 20), ReadPosRankSum outside the range −2 to 2, MQRankSum outside the range −2 to 2, and BaseQRankSum outside the range −2 to 2. For population genetic analyses, indels were removed using VCFtools v0.1.17[70], retaining only high-confidence SNPs. Only biallelic SNPs were retained and variants with ambiguous nucleotides were removed.

Functional annotation of variants

Functional annotation of variants relative to the reference genome annotation was performed using SnpEff v5.4[71]. Variants were classified according to their predicted effects on coding sequences (e.g. synonymous, missense, frameshift and stop-gained) and grouped into impact categories (HIGH, MODERATE, LOW and MODIFIER). To avoid redundant or ambiguous locus assignments, only the primary annotation reported by SnpEff—corresponding to the most severe predicted functional effect—was retained for each variant. Secondary annotations (e.g. upstream, downstream or intergenic effects) were discarded. Variant annotations were extracted from the annotated VCF using BCFtools v1.17[68]. For variants with multiple annotations, the primary annotation described above was retained. Variants with missing genotype calls were treated as missing data.

Gene presence–absence analysis

Gene presence/absence variation across isolates was assessed by calculating sequencing read coverage across gene coordinates using BEDTools v2.31.1[72]. Gene coordinates were obtained from the reference genome annotation (GFF file), and coverage was computed from BAM files for each isolate. Genes with ≥10% of their length covered by reads were classified as present, whereas genes below this threshold were considered absent. The resulting gene-by-isolate matrix was used to investigate gene loss and accessory genome variation. Custom scripts implemented in Bash, AWK and R (tidyverse) were used for parsing, data aggregation and generation of intermediate datasets.

Gene-level mutation matrices and functional status

To quantify coding variation, gene-by-isolate matrices were constructed describing: (i) gene presence/absence; (ii) occurrence of high-impact mutations (e.g. frameshift and stop-gained); (iii) synonymous substitutions and; (iv) missense mutation counts. Protein lengths were extracted from SnpEff annotations based on amino acid position and length information associated with coding variants. These values were used to calculate a normalized protein conservation score for each gene and isolate.

All matrices were integrated into a unified gene-by-isolate functional status matrix using a hierarchical classification framework. Genes that were absent or carried at least one HIGH-impact mutation were classified as non-functional (0). Genes containing missense variants were assigned their corresponding protein conservation score. Genes present without missense mutations, or containing only synonymous substitutions, were classified as intact (value=1). This approach prioritizes gene loss and disruptive mutations over lower-impact variation.

Population network and structure

Phylogenetic networks were inferred using SplitsTree v6.7.5[73]. Population structure was initially explored using principal component analysis (PCA) implemented in PLINK[68]. The first two principal components were visualized to assess major axes of genetic differentiation among samples. PCA plots were generated in R using the ggplot2 package. Population genetic structure was further inferred using the variational Bayesian framework implemented in fastStructure v.0.0.0[74]. Analyses were performed using the PLINK binary dataset as input for values of *K* ranging from 1 to 10 in order to explore alternative numbers of genetic clusters. For each value of *K*, fastStructure was run using default parameters. The optimal number of clusters was determined using the chooseK.py utility based on model complexity and marginal likelihood, and was independently supported by cross-validation analysis implemented in ADMIXTURE v1.3.0[75]. Individual ancestry coefficients were visualized as bar plots in R using sample information from PLINK .fam files.

**Disruption of the selected gene**

Targeted deletion of the PKS–NRPS gene (*CLUP02_01846*) in *C. lupini* was achieved by homologous recombination using *Agrobacterium tumefaciens*-mediated transformation (ATMT)[76,77]. Briefly, ~1–1.5 kb genomic regions upstream and downstream of the *CLUP02_01846* coding sequence were amplified from genomic DNA and cloned into a binary vector flanking a hygromycin B phosphotransferase resistance cassette, thereby generating the deletion construct. Correct assembly of the plasmids was confirmed by restriction enzyme digestion and Sanger sequencing using universal primers (M13F/M13R) together with gene-specific primers designed to span the recombination regions through a primer-walking strategy.

Validated constructs were introduced into competent *A. tumefaciens* cells and used for transformation of *C. lupini* conidia following established ATMT protocols. After co-cultivation, transformants were selected on hygromycin B-containing medium supplemented with antibiotics to counterselect *A. tumefaciens*. Emerging colonies were single-spore subcultured to obtain stable transformants and were subsequently subjected to molecular screening.

Primary screening was performed by PCR using primer pairs positioned outside the homologous flanking regions to verify correct gene replacement by homologous recombination. Additional PCR assays targeting internal regions of *CLUP02_01846* were used to confirm loss of the native locus and to exclude retention of the wild-type allele.

To validate gene deletion at the genomic level, whole-genome sequencing was performed on selected transformants. Genomic reads were assembled using SPAdes v3.15.1, and the resulting assemblies were used as local databases to confirm both the absence of *CLUP02_01846* and the presence of the hygromycin resistance cassette at the target locus. Selection of transformants were performed as previously described[78]. Two independent deletion mutants, Δ*CLUP02_01846*α and Δ*CLUP02_01846*β, were selected and used for subsequent phenotypic and pathogenicity assays.
